# High-Resolution Multimodal Profiling of Human Epileptic Brain Activity via Explanted Depth Electrodes

**DOI:** 10.1101/2024.07.03.601925

**Authors:** Anuj Kumar Dwiwedi, Arun Mahesh, Albert Sanfeliu, Julian Larkin, Rebecca Siwicki, Kieron J. Sweeney, Donncha F. O’Brien, Peter Widdess-Walsh, Simone Picelli, David C. Henshall, Vijay K. Tiwari

## Abstract

Understanding neurological disorders necessitates systems-level approaches that integrate multimodal data, but progress has been hindered by limited sample availability, and the absence of combined electrophysiological and molecular data from live patients. Here, we demonstrate that intracranial stereoelectroencephalography (SEEG) electrodes used for identifying the seizure focus in epilepsy patients enable the integration of RNA sequencing, genomic variants and epigenome maps with in vivo recordings and brain imaging. Specifically, we report a method, MoPEDE (Multimodal Profiling of Epileptic Brain Activity via Explanted Depth Electrodes) that recovers extensive protein-coding transcripts, DNA methylation and mutation profiles from explanted SEEG electrodes matched with electrophysiological and radiological data allowing for high-resolution reconstructions of brain structure and function in human patients. Our study shows that epilepsies of different aetiologies have distinct molecular landscapes and identify transcripts correlating with neurophysiological signals, including immediate early genes, inflammation markers, and axon guidance molecules. Additionally, we identify DNA methylation profiles indicative of transcriptionally permissive or restrictive chromatin states. While gene expression gradients corresponded with the assigned epileptogenicity index, we found outlier molecular fingerprints in some electrodes, potentially indicating seizure generation or propagation zones not detected during electroclinical assessments. These findings validate that RNA profiles, genetic variation and genome-wide epigenetic data from explanted SEEG electrodes offer high-resolution surrogate molecular landscapes of brain activity. Our transformative MoPEDE approach has the potential to enhance diagnostic decisions and deepen our understanding of epileptogenic network processes in the human brain.

## INTRODUCTION

An improved understanding of human brain function requires systems-level approaches that sample and integrate multimodal data spanning several orders of magnitude from single molecules, through individual cells to local networks and up to higher order organisation.^1, 2^ Combining RNA sequencing and genome architecture maps with *in vivo* recording technologies and brain imaging enables the production of high-resolution reconstructions of the structure and function of the mammalian brain in health and disease. Indeed, increasingly comprehensive taxonomies of the mammalian brain have been achieved by integrating sequencing approaches with other modalities. These studies have demonstrated neuronal phenotypes including morphology, location and electrophysiologic properties are strongly defined by cell type-specific transcriptional and epigenetic signatures for different neuron subtypes,^3–5^ across different layers of the cortex,^3, 4, 6^ and structures throughout the mouse brain including the hippocampus.^5–7^ Transcription factors are a principle driver of cellular diversity, regional specialisation of function and neurotransmitter type which in turn establishes the electrophysiological properties of different cell types.^7^ Equivalent mapping exercises for the human brain have emerged, with recent single-cell transcriptional and epigenetic studies revealing the distinct molecular programs that define neuronal and non-neuronal cell type and diversity, network and regional organisation, and complexity.^8–10^ These insights utilize, however, postmortem tissue and access to living brain tissue is a key factor for functional characterisation of the human brain.

Treatment-resistant epilepsies represent unique opportunities for access to the living human brain. *Ex vivo* electrophysiologic, pharmacologic and gene expression profiling of surgically-resected material has enabled important advances into brain function in health and disease.^11, 12^ The analysis from implanted electrodes, placed to guide surgical decisions has also enabled studies of human brain function^13^, including elucidating network behaviour during seizure onset,^14^ and how epileptiform activity can interfere with memory.^15^ Intracranial electroencephalography (EEG) recordings from stereotactically-implanted electrodes (known as stereoelectroencephalography or SEEG) are performed for a subset of patients with difficult-to-localize focal epilepsy to identify seizure-onset zones.^16^ The implantation of typically between 5 – 15 electrodes with multiple contact sites into deep brain structures yields necessary spatial and temporal mapping of hyperexcitable epileptogenic tissue, enabling localization of the seizure onset zone, the associated propagation zones and differentiation from the normal brain.^13, 16^ The neurophysiologic findings are then combined with imaging, computational tools and clinical information to guide surgical decisions, with operations performed upon explantation of SEEG electrodes.^16–18^ Researchers recently reported that DNA present on explanted SEEG electrodes can be used to identify somatic mutations in pre-resection epilepsy patients.^19–21^ Notably, a gradient of mosaic gene variation may be present in relation to recorded epileptiform activity.^20^ Mapping epigenetic marks using such material may also have utility for subtyping malformations of cortical development such as focal cortical dysplasias.^22^ Assembling comprehensive molecular architectures that comprise read-outs of gene activity, for example, seizure-regulated gene transcripts or inflammatory signals, in combination with recorded neurophysiology would offer powerful new insights into human brain function and causal mechanisms of epilepsy, and potentially support surgical decision-making. There are, however, several unknowns. What additional nucleic acids can be reliably obtained from explanted electrodes, in particular mRNA transcripts, and do these correspond to epigenetic marks that influence chromatin state? Do the signals retain information about cell types and implantation locations? Finally, epilepsies are heterogeneous in clinical presentation, semiology and underlying mechanism so can this approach be applied across surgical candidates with different aetiologies and is it scalable from highly focal epilepsies through to those affecting larger structures and networks? Indeed, defining the extent of the epilepsy is one of the greatest challenges in epileptology because of what we cannot ‘see’ with existing technology and hence, novel methodological developments are urgently required.

Here we report a method called MoPEDE (Multimodal Profiling of Epileptic Brain Activity via Explanted Depth Electrodes) in which an extensive repertoire of protein-coding transcripts, DNA methylation and variant profiles can be recovered from explanted SEEG electrodes matched with electrophysiological and radiological data. We demonstrate proof-of-concept that epilepsies of distinct aetiology are associated with distinct molecular landscapes and identify transcripts whose levels are tracked with recorded neurophysiological signals. This included functional categories such as immediate early genes, markers of inflammation and axon guidance. We identify DNA methylation profiles consistent with transcriptionally permissive or restrictive chromatin states. Notably, while gradients of gene expression between sites corresponded to the assigned level of epileptogenicity, we found outlier electrode molecular fingerprints which could indicate seizure spread or generation zones not assigned during clinical neurophysiologic assessment. Together, these findings provide proof-of-concept that RNA profiles, genome-wide epigenetic surveillance and mutation analysis can be obtained from explanted SEEG electrodes and these provide surrogate molecular landscapes of human brain activity at high resolution that may support diagnostic decisions as well as improve our understanding of the epileptogenic process in the human brain.

## RESULTS

### Multimodal profiling of different epilepsy subtypes from SEEG electrodes

The SEEG method utilizes intracranial electrodes to measure electrophysiological activity in various brain regions, including deeper structures such as the hippocampus, amygdala, and insula, providing a comprehensive sampling of these areas. Previous studies have demonstrated that these electrodes can be used to identify brain-specific somatic mutations in different epileptic brain regions.^19–21^

Here, we collected explanted SEEG electrodes from patients with three distinct epilepsy subtypes (see below). Based on EEG profiles, we stratified the electrodes into three major categories: 1) Seizure Onset Zone (SOZ), 2) Propagation Zone (PZ), and 3) Non-Involved Zone (NIZ). We then sectioned the electrodes from the SOZ, PZ, and NIZ for each patient and extracted RNA and genomic DNA (gDNA) from these samples, performing both transcriptome and epigenome (DNA methylation) profiling (Fig. 1A). We further performed variant analysis from the derived RNA profiles (Fig. 1A).

**Figure 1.**
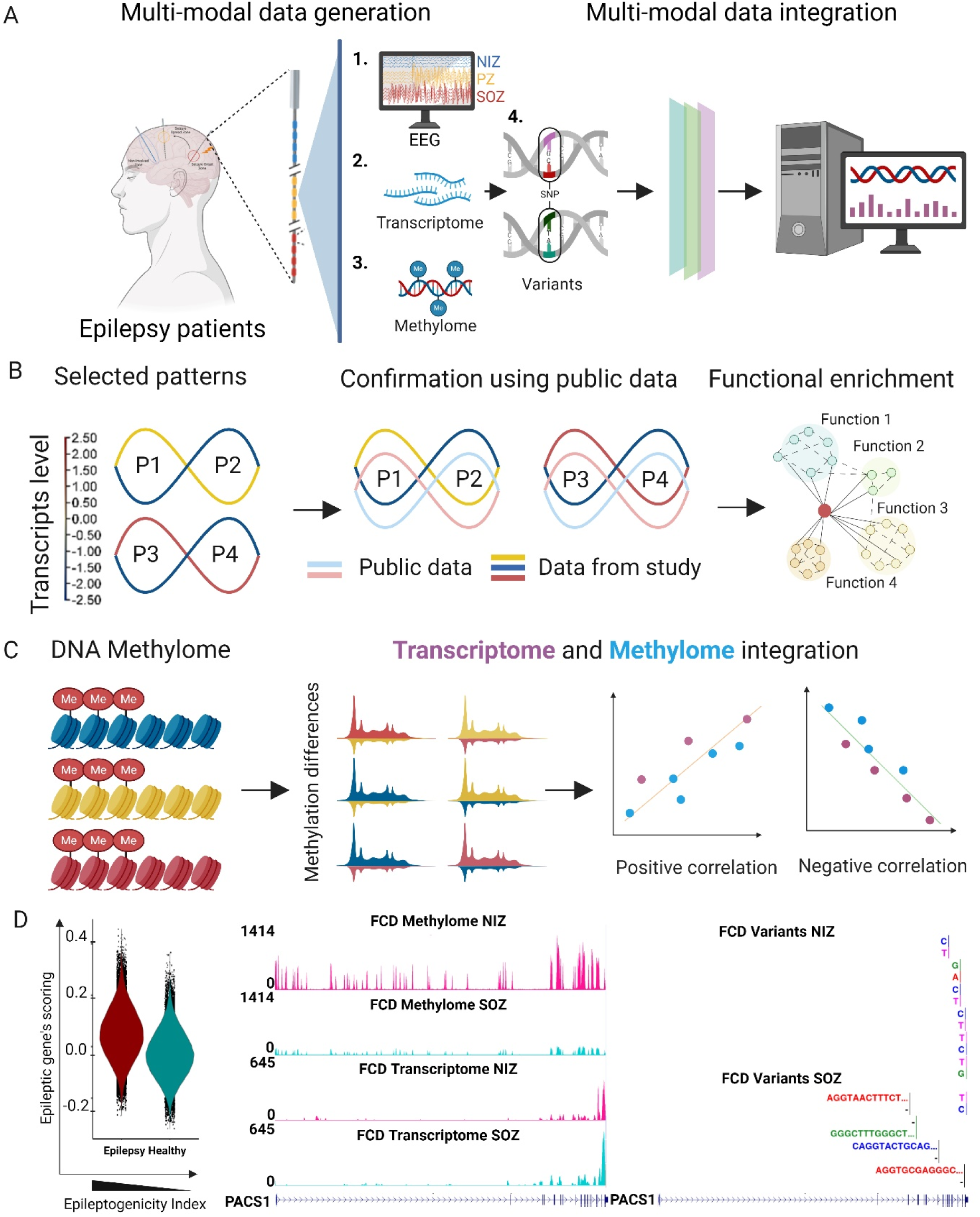
Multimodal profiling of different epilepsy subtypes using SEEG electrodes. (A) We generated single-source electroencephalogram (EEG), whole transcriptome, methylome and variants profiles from electrodes collected from focal cortical dysplasia (FCD), non-lesional temporal lobe epilepsy (TLE), and Rasmussen’s encephalitis (RE) brains. (B) Multimodal data integration was applied to identify molecular signatures and disease causes. We distinguished different sets of signatures based on K-means clustering and validated their patterns using publicly available epilepsy data, followed by functional enrichment analysis. (C) Whole-methylome profiles were generated using the same samples and identified differentially methylated regions by investigating positive and negative correlations between the transcriptome and methylome data. (D) Panel shows a snapshot of the integration of electrophysiology data with transcriptome signatures and a single-resolution map illustrating the correlation between transcriptome ,methylome and variants levels for a known epilepsy risk-associated gene (*PACS1*) in both onset and cold electrodes from an FCD brain.

We first analyzed the transcriptome data, focusing on specific expression patterns (P) between the SOZ, PZ, and NIZ (Fig. 1B). These were designated as Pattern 1 (P1): Transcripts with higher expression in the PZ and lower levels in the NIZ. Pattern 2 (P2): Transcripts with higher expression in the NIZ and lower expression in the PZ. Pattern 3 (P3): Transcripts with higher expression in the SOZ and lower expression in the NIZ. Pattern 4 (P4): Transcripts with higher expression in the NIZ and lower expression in the SOZ. We compared these patterns (P1-P4) with a publicly available epilepsy dataset,^23^ and conducted functional enrichment analysis to identify significant biological pathways (Fig. 1B). For DNA methylation analysis, we examined the same samples (SOZ, PZ, and NIZ) from the three patients to identify differentially methylated regions (DMRs). We integrated the transcriptome data with the methylome data to understand how DNA methylation profiles correlate with corresponding transcriptome profiles and disease etiology (Fig. 1C).

Finally, we aimed to integrate the SEEG neurophysiology (visual analysis and epileptogenicity index score) with our transcriptome and DNA methylome data as well as variants called using the RNA-seq profiles (Fig. 1D). This multimodal approach has the potential to demonstrate how EEG-based epileptogenicity index scores, RNA, and gDNA from SEEG electrodes of patients with epilepsy can be utilized to profile and understand the molecular underpinnings of epilepsy. This may ultimately lead to a personalized molecular diagnosis and more targeted and effective treatments.

### Patient neurophysiology, neuroimaging, and neuropathological features

To assess the applicability of our method for the different epilepsy subtypes, we recruited three patients with epileptic foci of diverse aetiology and extent: focal cortical dysplasia (FCD), non-lesional temporal lobe epilepsy (TLE), and Rasmussen’s encephalitis (RE). All participants underwent robot-assisted SEEG monitoring. Depth electrodes with twelve, fifteen, or eighteen contacts were implanted based on a pre-operative hypothesis of the SOZ. By standard nomenclature, each depth is labelled with a letter, with electrode contacts numbered from mesial to lateral. The anatomic location of each electrode was confirmed by postoperative computed tomography (CT) co-registered with preoperative volumetric magnetic resonance imaging (MRI) (Figure 2). Continuous EEG recordings of seizures was conducted with concurrent video, and visual analysis classified cortical regions into the SOZ, PZ, or NIZ. For quantitative SEEG analysis, representative seizures with minimal artefacts occurring over 48 hours post-implantation were selected. The epileptogenicity index (EI) at each electrode pair was measured to assess changes in energy ratio and time delay from electrode contacts from seizure onset, estimating the epileptogenicity of cortical regions.^24, 25^ We then analysed the biological samples to identify molecular characteristics and compared them across the patient groups.

**Figure 2.**
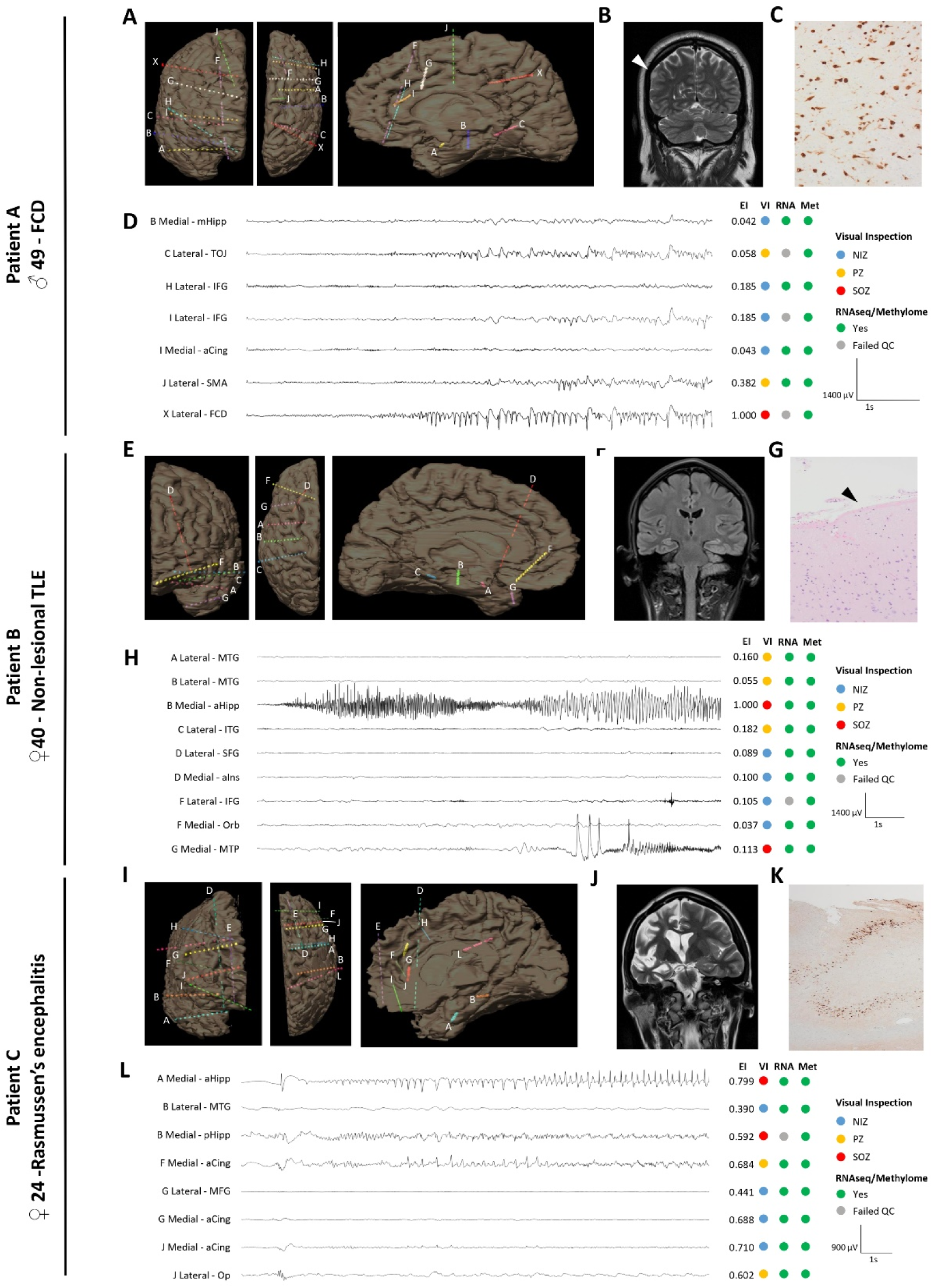
Neurophysiology, neuroimaging, and neuropathological features of study participants. (A, E, I) Coronal, axial, and mesial sagittal volumetric MRI reconstruction showing the location of the implanted SEEG electrodes for patients A, B, and C. (B) Coronal T2-weighted MRI sequence of patient A. Arrow indicates the location of the FCD. (C) NeuN immunohistochemical staining for patient A demonstrated dysmorphic, haphazardly arranged and abnormally clustered cortical neurons consistent with FCD, ILAE Type IIa. Balloon cells were not identified. (D, H, L) Representative sample SEEG ictal recording with matched Epileptogenicity Index, epileptogenic network involvement based on visual inspection (VI) and presence or absence of contact points in our molecular analysis. Letters correspond to depth electrodes indicated in A, E, and I, bipolar recording. At ictal onset, there is fast repetitive spiking and paroxysmal fast activity. High pass filter 5Hz, low pass filter 80Hz, Notch 50Hz RNAseq: RNA sequencing; Met: Methylome sequencing; NIZ: Non-involved Zone, PZ: Propagation Zone, SOZ: Seizure Onset Zone, MTG: middle temporal gyrus, mHipp: mid-hippocampus, TOJ: temporo-occipital junction, pHipp: posterior hippocampus, aHipp: anterior hippocampus, SFG: superior frontal gyrus, MFG: middle frontal gyrus, IFG: inferior frontal gyrus, Orb: orbitofrontal, aCing: anterior cingulate, SMA: supplementary motor area, ITG: inferior temporal gyrus, aIns: anterior insula, MTP: mesial temporal pole, Op: opercular. (F) Coronal MRI FLAIR sequence of patient B demonstrating normal temporal lobe structures. (G) Haematoxylin and eosin staining for patient B showing Chaslin’s subpial gliosis, indicated by arrow. (J) Coronal T2-weighted MRI sequence of patient C showing diffuse right hemisphere atrophy including the hippocampus, consistent with RE. (K) NeuN immunohistochemical staining for patient C showing severe hippocampal sclerosis with granular neuron depletion, dispersion and mossy fibre sprouting. No evidence of active encephalitis was present.

The first patient was a 49-year-old male with medication-resistant focal epilepsy due to a right parietal FCD type IIA (Fig. 2A-D). During SEEG monitoring, thirty-three electroclinical seizures were recorded, with EEG onset from the lateral contacts of the depth electrodes at the parietal FCD (Table 1). Seizure propagation involved the supplementary motor area and temporo-occipital junction. The patient underwent resection of the FCD and achieved an Engel class 1A outcome, remaining seizure-free at one year. Histopathology confirmed a type IIa FCD. We collected 18 electrode samples from NIZ, PZ and SOZ regions and then selected ∼40% of electrodes covering the NIZ, PZ and SOZ regions for whole transcriptome and DNA methylome profiling (Fig. 2D).

**Table 1:** Clinical characteristics of patients.

|  | Patient A | Patient B | Patient C |
| --- | --- | --- | --- |
| **Age (years)** | 49 | 40 | 24 |
| **Sex** | M | F | F |
| **Age at epilepsy onset (years)** | 15 | 32 | 15 |
| **Anti-seizure medications (prev. ASMs)** | CAR, LEV, TOP<br>(BRIV, GAB, LAC, LTG, PHEN, VAL, VIG) | CEN, ESL<br>(BRIV, CLOB, CLON, LAC, LTG) | CLOB, LEV, PRE, ZON<br>(CEN, CLON, ESL, LAC, LTG, PER, ZON, TOP) |
| **Neuroimaging** | FCD | Non-lesional | Diffuse right hemi-atrophy, gliosis |
| **Duration of SEEG monitoring (days)** | 12 | 18 | 11 |
| **No. of depths** | 9 | 6 | 10 |
| **Surgery** | Lesionectomy | Left ATL with AH | Right ATL with AH, inferior frontal resection with prefrontal disconnection |
| **Histopathological diagnosis** | FCD IIa | Chaslin's subpial gliosis | Hippocampal sclerosis, cortical gliosis, no inflammation |
| **Surgical outcome, Engel Class** | 1A | 2A | 2A |

*FCD: focal cortical dysplasia, ATL: anterior temporal lobectomy, AH: amygdalo-hippocampectomy, CAR: carbamazepine, LEV: levetiracetam, TOP: topiramate, BRIV: brivaracetam, GAB: gabapentin, LAC: lacosamide, LTG: lamotrigine, PHEN: phenytoin, VAL: valproate, VIG: vigabatrin, CEN: cenobamate, ESL: eslicarbazepine, CLOB: clobazam, CLON, clonazepam, ZON: zonisamide, PRE: pregabalin, PER: perampanel.*

The second patient was a 40-year-old female with MRI-negative, medication-resistant left TLE (Fig. 2E-H and Table 1). SEEG monitoring recorded six electroclinical seizures, with EEG onset from the anterior hippocampus and the temporal pole. Seizure propagation involved the middle temporal gyrus and inferior temporal gyrus. The patient underwent a left temporal lobectomy with amygdalo-hippocampectomy, resulting in an Engel class 2A outcome at one year, initially seizure-free but experiencing rare seizures subsequently. Histopathology revealed Chaslin’s subpial gliosis. We collected 12 electrode samples from the NIZ, PZ, and SOZ regions. We then profiled approximately 75% of these electrodes, covering all three regions (NIZ, PZ, and SOZ), for whole transcriptome and DNA methylation (Fig. 2H).

The third patient was a 24-year-old female diagnosed with RE at age 15 years, treated medically (Fig. 2I-L). Persistent medication-resistant epilepsy led to SEEG monitoring, during which 32 electroclinical seizures were recorded, primarily from the hippocampus and the gyrus rectus. Seizure propagation involved the cingulate and frontal operculum. The patient underwent right frontal and anterior temporal resection with an Engel class 2A outcome at one year, initially seizure-free with rare seizures subsequently. Initial brain biopsy at age 15 years showed chronic encephalitis with T-cell rich perivascular and parenchymal inflammation. Histopathology from the resection eight years later revealed severe hippocampal sclerosis and cortical gliosis, with no active inflammation. We collected 20 electrode samples from the NIZ, PZ, and SOZ regions. We then profiled all three regions (NIZ, PZ, and SOZ), encompassing approximately 40% of the electrodes, for whole transcriptome and DNA methylation (Fig. 2L).

After we collected the explanted SEEG contacts from these three patients, we first investigated whether these retained any intact cells. For this, electrodes were cut into small pieces and rinsed with PBS for collecting cells. Trypan blue staining confirmed the presence of cells on these electrodes (Supplementary Fig. 1A). We then extracted total nucleic acids (both RNA and gDNA) from these cells and measured their quantity using a Nanodrop spectrophotometer. Remarkably, we obtained significant concentrations of nucleic acids from these electrodes, with amounts directly proportional to the number of metal contact points in the sample (Supplementary Fig. 1B). Next, we analyzed the quality of these nucleic acids using a high-sensitivity fragment analyzer, finding both gDNA and RNA in considerable concentrations (Supplementary Fig. 1C-E). We then divided these samples into equal portions and purified the gDNA and RNA separately (Supplementary Fig. 1F and G). The purified RNA exhibited a acceptable RNA integrity numbers (RIN ranging from 2.5 to 8.4) (Supplementary Fig. 1G). These results indicate that SEEG electrodes used in pre-surgical evaluation of epilepsy carry sufficient cells from the implanted brain regions to enable transcriptome and epigenome of living people with epilepsy.

### Single-sourced multi-ome profiles of SEEG electrode contacts show high sequence coverage and mapping

To achieve comprehensive sequence coverage in the transcriptome of very low input RNA samples from the SEEG electrodes, we applied the Flash-seq method.^26^ Originally developed for full-length single-cell RNA sequencing, we adapted Flash-seq for bulk RNA sequencing in our study.^27^ This allowed us to obtain full-length transcripts with high coverage from the three patients (FCD, TLE, and RE) using electrodes representing the SOZ, PZ, and NIZ, performing both transcriptome and DNA methylome analysis (Fig 3A).

**Figure 3.**
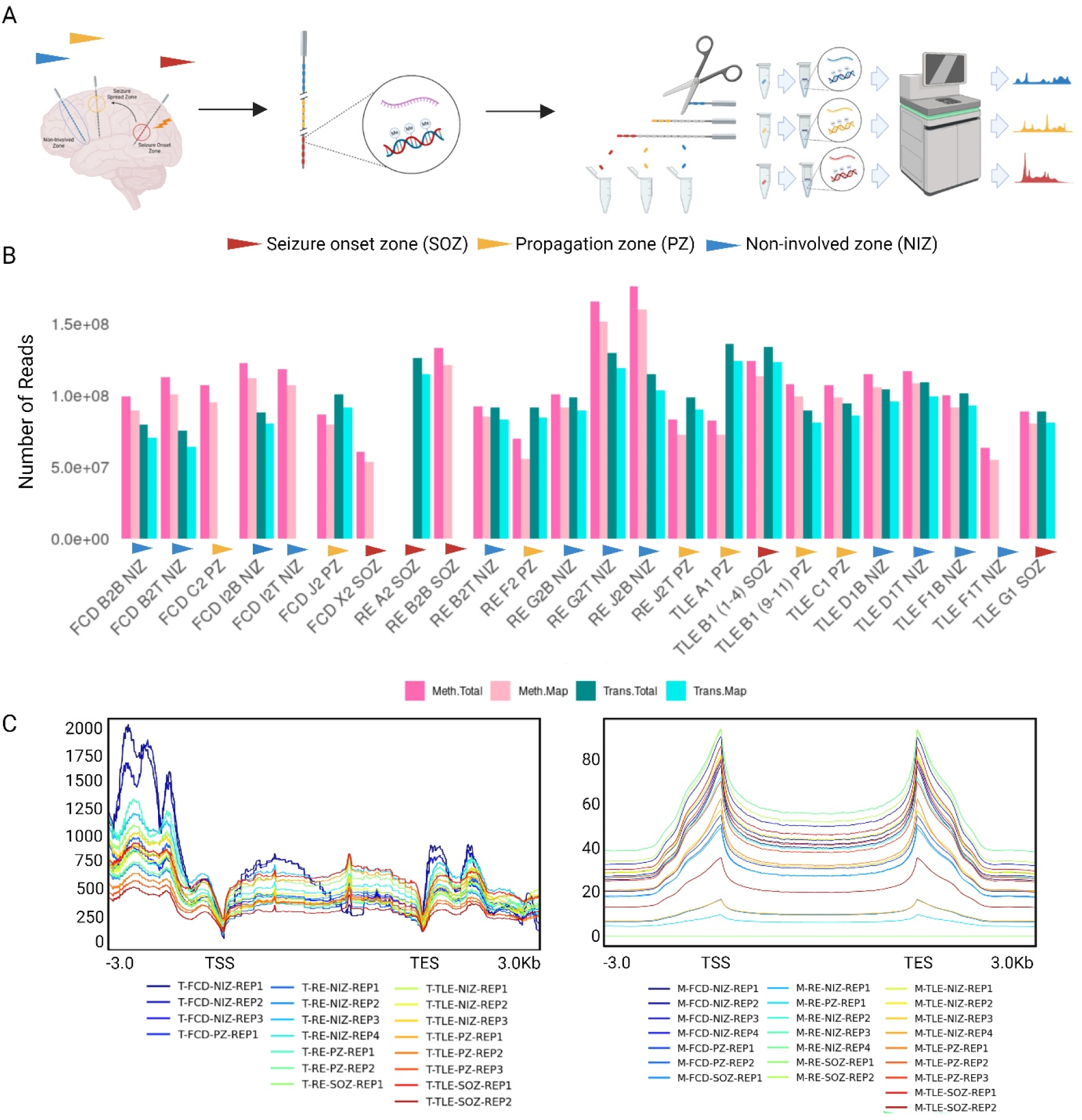
Single-source high throughput multi-omics (transcriptome and methylome) profile of EEG electrodes in epilepsy. (A) Selection of EEG electrodes: EEG electrodes were selected from three zones: seizure onset zone (SOZ), propagation zone (PZ), and non-involved zone (NIZ). The attached nucleic acid material was extracted from each electrode, and whole transcriptome and methylome data were generated using this same material. (B) Seven electrodes explanted from the brain of a patient with FCD were used. An average of 100 million and 85 million reads were sequenced for the methylome and transcriptome, respectively. Notably, 91 million and 74 million methylome and transcriptome reads, respectively, mapped to the human genome. Similarly, nine electrodes explanted from the brain of a patient with non-lesional TLE were used. An average of 100 million and 95 million reads were sequenced for the methylome and transcriptome, respectively. Prominently, an average of 95 million and 87 million methylome and transcriptome reads, respectively, mapped to the human genome. Additionally, eight electrodes explanted from the brain of a patient with REwere used. An average of 117 million and 101 million reads were sequenced for the methylome and transcriptome, respectively. Importantly, an average of 105 million and 92 million methylome and transcriptome reads, respectively, mapped to the human genome. (C) Transcriptome and methylome density: The transcriptome and methylome density of each electrode from all three disorders were plotted across gene body and flanking regions (3kb). A consistent negative correlation between methylation and transcription was observed across all electrodes.

From the FCD case, we used seven different electrode samples, achieving an average of 100 million and 85 million reads per sample in DNA methylome and transcriptome sequencing, respectively. Of these, 91 million reads from the methylome and 76 million reads from the transcriptome were mapped to the human genome (Fig. 2D, 3B). From the TLE case, we used nine electrode samples, resulting in an average of 100 million and 106 million reads per sample in DNA methylome and transcriptome sequencing, respectively. Some 90 million methylome reads and 97 million transcriptome reads were mapped for the TLE case to the human genome (Fig. 3B). One of the RNA samples failed in QC and was excluded from sequencing (Fig. 2H, 3B). From the RE case, eight electrode samples were analysed, yielding an average of 117 million and 101 million reads per sample in DNA methylome and transcriptome sequencing, respectively. Of these, 105 million methylome reads and 97 million transcriptome reads were mapped to the human genome (Fig. 3B). One of the RNA samples failed in QC and was excluded from sequencing (Fig. 3B).

We then performed density analysis for transcriptome reads across the transcriptional start sites (TSS), covering a 3kb upstream and downstream window (Fig. 3C left panel). Similarly, we analysed the density of DNA methylation reads across the TSS sites with a 3kb window (Fig. 3C right panel). As expected, these density plots revealed a consistent negative correlation between the transcriptome and DNA methylome across all samples analysed (Figs. 3C). These data demonstrate that it is possible to obtain high coverage multiome profiles from a few SEEG electrode contacts from pre-surgical epilepsy patients.

### Transcriptome analysis of SEEG electrodes reveals distinct signatures of epileptic brain regions

To delineate transcriptional changes associated with different epilepsy subtypes, we performed a differential expression analysis comparing SOZ, PZ and NIZ (Fig. 1B and Fig. 4A). To validate the transcriptome profile from SEEG electrodes, we analyzed previously published single nuclei RNA sequencing (snRNAseq) data,^23, 28^ from surgically-obtained epilepsy samples and matched controls (Supplementary Figure 2). We integrated epilepsy and healthy single nuclei profiles and annotated different clusters using markers for various excitatory and inhibitory neurons (Supplementary Figure 2A and B).

**Figure 4.**
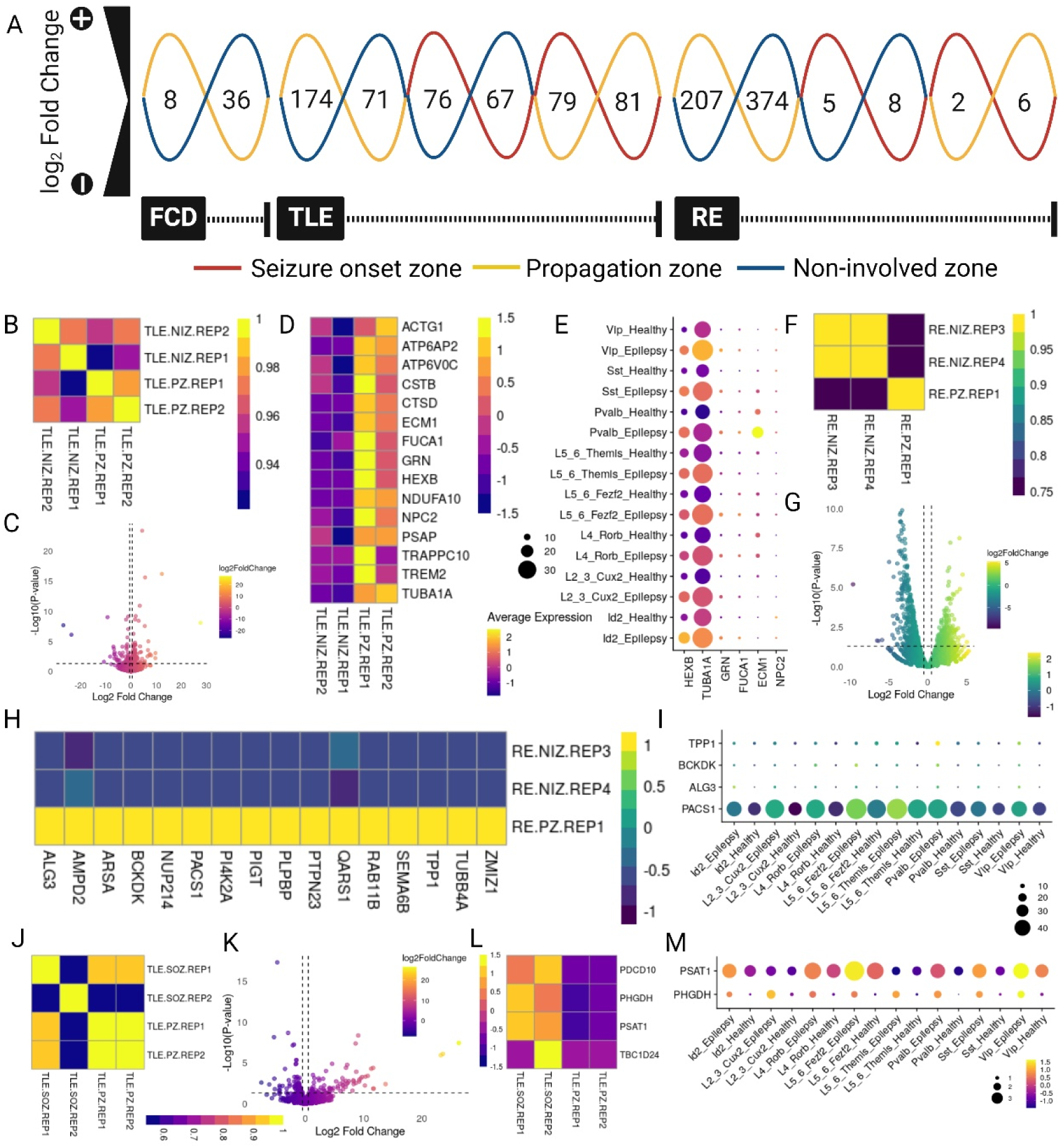
Transcriptional signatures activated in different epileptic brain regions. (A) Identification of transcriptional patterns in epileptic brains. The figure shows differentially expressed gene identified in three epilepsy subtypes: FCD, non-lesional TLE, and RE. . (B-M) Expression of known epilepsy associated genes in different brain regions in SOZ, NIZ, and PZ (where applicable) for each epilepsy subtype (TLE in (B-E and J-M), and RE in (F-I)). The data is presented alongside corresponding scRNA epilepsy data for comparison.

For the FCD patient sample, only the PZ and NIZ samples were sequenced. We performed replicate-wise transcriptome correlations comparing PZ vs NIZ and found they were poorly correlated (Supplementary Fig. 3A). We identified 8 significantly upregulated and 36 downregulated genes in the PZ compared to the NIZ (Fig. 4A, Supplementary Fig. 2B and Supplementary table 1 & 2). GO enrichment analysis indicated that genes activated in PZ areas were involved in translational processes (Supplementary Figure 2C) whereas the downregulated genes were mostly involved in metabolic processes (Supplementary Figure 2D). We next compared the differentially expressed genes with the publicly available epileptic-associated genes^28^ finding 2 genes overlapped, but both showed lower expression in the PZ compared to the NIZ (Supplementary Figure 2E). In contrast, we found genes that are downregulated in the PZ also showed downregulation in epilepsy samples in the public data set, including *RPL4, MTPAP* and *SNX30* (Supplementary Figure 2F).

For the TLE patient, we identified 174 significantly upregulated and 71 downregulated genes in the PZ compared to NIZ (Fig. 4A and 4C, and Supplementary table 3 & 4). The upregulated genes were enriched for translation, gene expression and peptide, and macromolecule biosynthetic pathways (Supplementary Figure 4A), while the downregulated genes were enriched for functions including superoxide generation, cytokine production and various stress signalling pathways (Supplementary Figure 4B). We then performed a correlation analysis between PZ and NIZ replicates and found they were anticorrelated, whereas a high correlation was observed within the replicates (Fig. 4B). When we compared the expression of these genes with epilepsy risk-associated genes^28^, we observed that transcripts enriched in the PZ sample contained more epilepsy-associated genes compared to NIZ (Fig. 4D and 4E).

Next, we compared the transcriptome of SOZ vs NIZ in TLE, finding there were 76 genes significantly upregulated and 67 genes downregulated, respectively (Fig. 4A, Supplementary Figure 4D and Supplementary table 4 & 6). A correlation analysis for SOZ vs NIZ found they are highly correlated (Supplementary Figure 4C). Epilepsy-associated genes were also highly expressed among the upregulated genes (Supplementary Figure 4E) and these genes were highly enriched for cellular responses to zinc, copper and cadmium ions (Supplementary Figure 4F), whereas the downregulated genes were enriched for immune response, cell junction disassembly and synapse pruning (Supplementary Figure 4G).

We further compared the transcriptome of SOZ vs PZ for TLE: 79 genes were significantly upregulated and 81 genes were downregulated, respectively (Fig. 4A, 4K and Supplementary table 7 & 8). The upregulated genes were enriched for cellular responses to zinc, copper and cadmium ions, dephosphorylation, cell fate commitment and stem cell differentiation (Supplementary Figure 5A), whereas the downregulated genes were enriched for proteolysis, protein catabolic process and peptidase activity (Supplementary Figure 5B). We also observed that epilepsy-associated genes *PDCC10, PHGDH, PSAT1* and *TBC1D24* were expressed at higher levels in SOZ compared to PZ (Fig 4L-M).

From the RE patient SEEG samples, we detected 207 significantly upregulated and 374 downregulated genes in the PZ compared to the NIZ (Fig. 4A, 4G and Supplementary table 9 & 10). When performing the correlation analysis, we found PZ vs NIZ were less correlated (Fig. 4F). Epilepsy associated genes (*TPP1, BCKDK, ALG3* and *PACS1*) were upregulated in PZ and highly expressed in epilepsy as compared with healthy control (Fig. 4I). The upregulated genes were associated with processes including nuclear export, amino acid transport, cell migration and positive regulation of gene expression (Supplementary Figure 6A), whereas the downregulated genes were enriched for translation, peptide and macromolecule biosynthesis and gene expression (Supplementary Figure 6B).

When we compared SOZ vs PZ in the RE patient we found 2 and 6 genes were significantly up and downregulated, respectively (Fig. 4A, Supplementary Figure 5D and Supplementary table 13 & 14). The upregulated genes were associated with the negative regulation of the apoptotic process, regulation of necrotic cell death and cellular respiration (Supplementary Figure 5E), whereas the downregulated genes were associated with the positive regulation of amyloid-beta formation, catabolic process and ubiquitin-dependent proteolysis (Supplementary Figure 5F).

In the SOZ compared to the NIZ, 5 genes were upregulated, and 8 genes were downregulated significantly (Fig. 4A and Supplementary Figure 6D) and they showed less correlation with each other (Supplementary Figure 6C). The upregulated genes in the SOZ were also highly expressed in epilepsy compared to healthy controls (Supplementary Figure 6F). GO enrichment analysis revealed that genes activated in the SOZ were involved in the negative regulation of cell cycle and cellular response to stress (Supplementary Figure 6E). In contrast, downregulated genes were associated with amyloid beta response and cholesterol import (Supplementary Figure 6G).

Overall, the SEEG transcriptional signatures from SOZ and PZ show a range of differential gene expression and shared as well as distinct biological processes for the three etiologies. Moreover, we successfully correlated our SEEG-recovered gene signatures with epilepsy scRNA-seq data from tissue-based transcriptomics, shedding light on the cell types harbouring these gene signatures and providing potential insights into the molecular and cellular mechanisms underlying different epilepsy subtypes.

### Integration of transcriptome and DNA methylome data provides insights into epigenetic dysregulation in epilepsy

We were then interested in studying variations in DNA methylation levels between the SOZ, PZ and NIZ across the three epilepsy subtypes. For FCD, this analysis showed that there is an overall raised levels of transcriptome and methylome in NIZ compared to PZ in the gene body and flanking regions and they are poorly correlated with each other (Supplementary Figure 7A, B and C). We then performed differential DNA methylation analysis between PZ and NIZ and found that 216 regions were hypermethylated and 1801 regions were hypomethylated (Supplementary Figure 7D). GO term analysis revealed that the hypomethylated regions were enriched for morphogenesis and GTPase mediated signal transduction (Supplementary Figure 7E), whereas hypermethylated genes were enriched for cholesterol storage and nuclear membrane disassembly (Supplementary Figure 7F).

For the TLE case we found there is an increase in methylation and transcriptome levels in NIZ compared to PZ and SOZ and they are less correlated with each other in PZ vs NIZ comparison (Fig. 5A, C, D, Supplementary Figure 9A and B). DMR analysis of these regions showed 2649 and 2405 hyper- and hypomethylated regions, respectively (Fig 5E).

**Figure 5.**
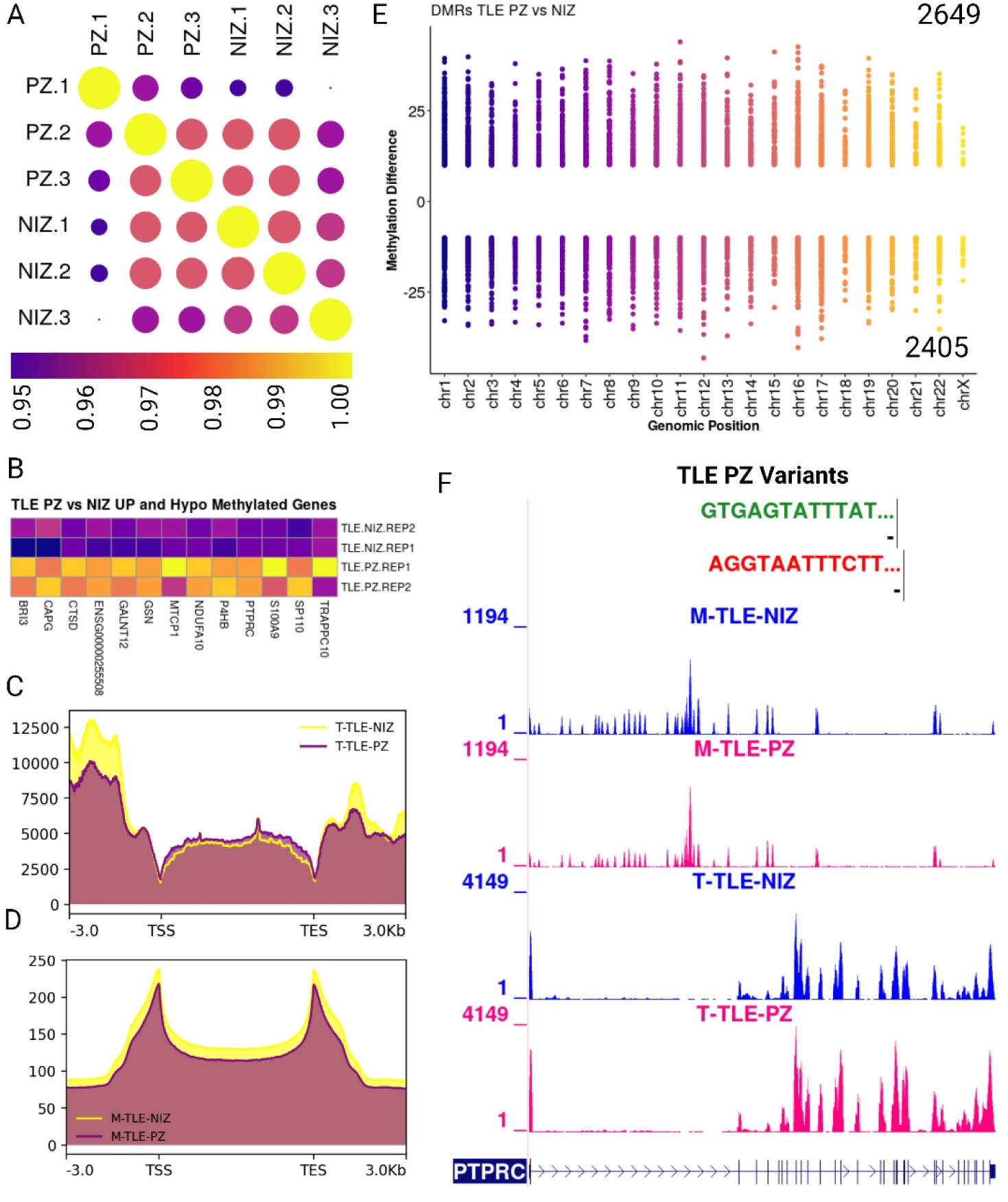
Multimodal single-base resolution map of epilepsy risk gene. (A) Replicate-wise correlation of methylation of CpG context from PZ and NIZ regions of TLE brain. (B and E) Differentially methylated regions in PZ vs NIZ comparison with genes negatively correlated with their transcriptome and methylome level. (C and D) Transcriptome and methylome levels are elevated in NIZ electrodes compared to PZ electrodes in TLE brain. Notably, transcriptome and methylome show an antagonistic correlation across the gene body. Decreased methylome levels in these regions accompany increased transcriptome levels in the gene body and flanking regions. Conversely, decreased transcriptome levels at the transcription start site (TSS) and transcription end site (TES) coincide with increased methylome levels at these regions. (E) A higher number of hypermethylated regions (2649) compared to hypomethylated regions (2405) were identified in PZ vs NIZ electrode comparisons. (F) Integration of transcriptome and methylome data revealed an inverse relationship between methylation and gene expression. A single-base resolution map was generated for *PTPRC* an epilepsy risk-associated gene, showing that depletion of methylation levels increased transcriptome levels in PZ electrodes. (F) Conversely, increased methylation levels correlated with decreased transcriptome levels in NIZ electrodes for *PTPRC*. In addition, we also observed PTPRC harboured in PZ regions in TLE brain.

We next investigated the link between gene expression and DNA methylation in PZ and NIZ regions in the TLE case by integrating single-source transcriptome and methylome data. This analysis revealed that methylation levels in PZ vs NIZ were negatively correlated with their transcriptome as expected (Fig. 5B and Supplementary Figure 8A). Notably, transcriptome and methylome showed an inverse correlation across the gene body and flanking regions (Fig. 5C and D). For example, decreased DNA methylation levels in these regions accompanied increased gene expression levels. Conversely, decreased transcription at the transcription start site (TSS) and transcription end site (TES) coincide with increased DNA methylation levels at these sites. GO term analysis revealed that the hypomethylated regions were enriched for phosphorylation and negative regulation of DNA binding of transcription factors (Supplementary Figure 8B), whereas the hypermethylated regions were enriched for GTPase-mediated signal transduction, regulation of cell migration and phagocytosis (Supplementary Figure 8C). Using the example of epilepsy-associated gene *PTPRC,* we found a reduction in the DNA methylation (hypomethylation) level and an increase in transcription in the PZ compared to NIZ (Fig. 5F).

Next, we analysed the correlation between the SOZ and NIZ DNA methylation levels in the TLE case and found less correlation between SOZ and NIZ (Supplementary Figure 8D). DMR analysis for these regions showed that 636 regions were hypermethylated and 475 regions were hypomethylated when comparing the SOZ to the NIZ area (Supplementary Figure 7E). GO-term analysis of hypomethylated regions was enriched for the adherence junction assembly, protein localisation to the cilium and negative regulation of transcription (Supplementary Figure 8F), whereas the hypermethylated regions were enriched for auditory behaviour, vocal learning and protein localisation to the membranes (Supplementary Figure 8G).

We found a positive correlation in DNA methylation levels among SOZ and PZ replicates (Supplementary Figure 9C). DMR analysis for these regions showed that 608 regions were hypermethylated and 872 regions were hypomethylated when comparing the SOZ to the PZ region (Supplementary Figure 8B). GTPase-mediated signal transduction, G-protein coupled receptor signalling and negative regulation of cell proliferation-related terms were enriched for the hypomethylated regions (Supplementary Figure 9E). The hypermethylated regions were enriched for processes including phospholipid biosynthesis and central nervous system development (Supplementary Figure 9F).

Tissue DNA methylation landscapes have recently been reported for RE^29^. For the SEEG samples from the RE patient, we observed elevated transcriptome and DNA methylome levels in NIZ as compared to PZ and SOZ regions in gene body and flanking regions (Supplementary Figure 10A and B). In addition, overall transcriptome and methylome levels were inversely proportional to each other in gene body and flanking regions. Next, DNA methylation correlation analysis showed a close correlation among PZ and NIZ replicates (Supplementary Figure 10C). DMR analysis for these regions showed that 31 regions were hypermethylated and 328 regions were hypomethylated when comparing the SOZ to the PZ region (Supplementary Figure 10D). Further, we observed transcriptionally upregulated genes negatively correlating with their DNA methylation levels (hypo-methylated) in PZ as compared with NIZ (Supplementary Figure 10E). GO-term analysis of hypomethylated regions was enriched for sulfate biosynthesis, mesenchymal cell migration and semaphorin-plexin signalling pathways (Supplementary Figure 10F).

### Combining neurophysiology with multiomic data provides deeper insights into epilepsy aetiology

Next, we explored the relationship between epileptogenicity index (EI) scores based on the recorded neurophysiologic signals and transcriptomic signatures derived from our differential gene expression analysis. To accomplish this, we integrated the EI scores of the TLE, and RE cases with transcriptome-derived signatures from NIZ, PZ, and SOZ and compared them with independent public epilepsy datasets.^23^ This analysis revealed that SOZ transcriptome signatures associated with high EI scores displayed elevated expression in epilepsy patient samples compared to healthy controls (left panel in Fig. 6A and B). Similarly, transcriptional signatures linked with moderate EI scores in the PZ of the TLE and RE cases showed increased activation in epilepsy samples relative to healthy controls (left panel in Fig. 6A).

**Figure 6.**
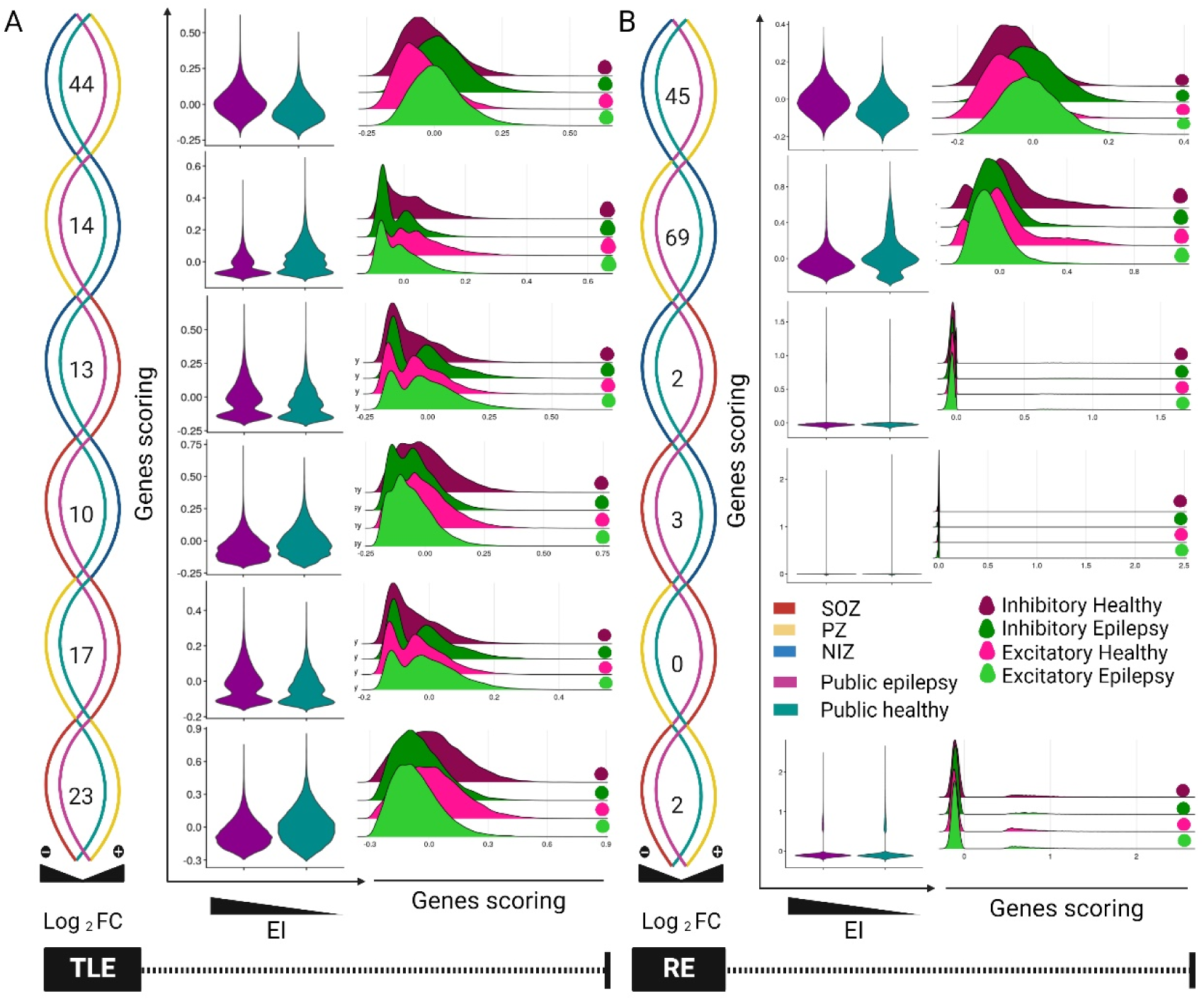
Transcriptional signatures associated with epileptogenicity index in epilepsy subtypes. (A and B) Relationship between the epileptogenicity index and transcriptional signatures in TLE and RE brains. This panel displays the relationship between the epileptogenicity index and transcriptional signatures across (set gene differentially expressed in the same direction in our and public data) different brain regions. We analyzed SOZ, PZ, and NIZ regions of non-lesional TLE and RE. The results show that transcriptional signatures associated with a high epileptogenicity index in TLE and RE were also highly activated in epilepsy samples compared to healthy controls (data from an independent public dataset). Similarly, transcriptional signatures associated with a moderate epileptogenicity index in non-lesional TLE and RE displayed increased activation in epilepsy compared to healthy controls (data from an independent public dataset). (A and B) The activity of transcriptional signatures in excitatory and inhibitory neurons. The ridge plots show the activity of transcriptional signatures within excitatory and inhibitory neurons. We compared the scores of signature genes associated with a high epileptogenicity index in excitatory and inhibitory neurons from both epilepsy and healthy samples. The results revealed higher correlation between activity scores and epileptogenicity index for these signature genes in excitatory and inhibitory neurons of epilepsy samples compared to their counterparts in healthy individuals.

Last, we explored the activity of these transcriptional signatures in excitatory and inhibitory neurons from both epilepsy and control samples. Our findings indicated that gene signatures from TLE, and RE were positively correlated with EI scores in both excitatory and inhibitory neurons of epilepsy patients compared to control counterparts (right panel Fig 6A-B). Overall, our analyses underscored that the expression profiles of gene signatures from SOZ and PZ of TLE, and RE patients closely mirrored those observed in existing epilepsy datasets in contrast to controls. This validation of our proof-of-concept study and the methodologies employed confirms the relevance of our approach in identifying and characterizing epilepsy-related transcriptomic signatures.

### Clinically significant genomic variants can be detected from RNA derived from SEEG contacts

Finally, we investigated whether genomic variants can be detected in the SEEG-derived RNA transcript sequences. Therefore we explored the presence of short variants (SNPs and Indels) in our SEEG-derived transcriptome data using GATK^30^ best practices workflows and annotated the high-quality variants through wANNOVAR (Fig. 7A). We then mapped the chromosome-wide distribution of variants in RE where we compared the difference between NIZ and PZ (Fig. 7B). Similarly, we performed a chromosome-wide distribution of variants in TLE where we compared the difference between NIZ, PZ and SOZ (Fig. 7C).

**Figure 7.**
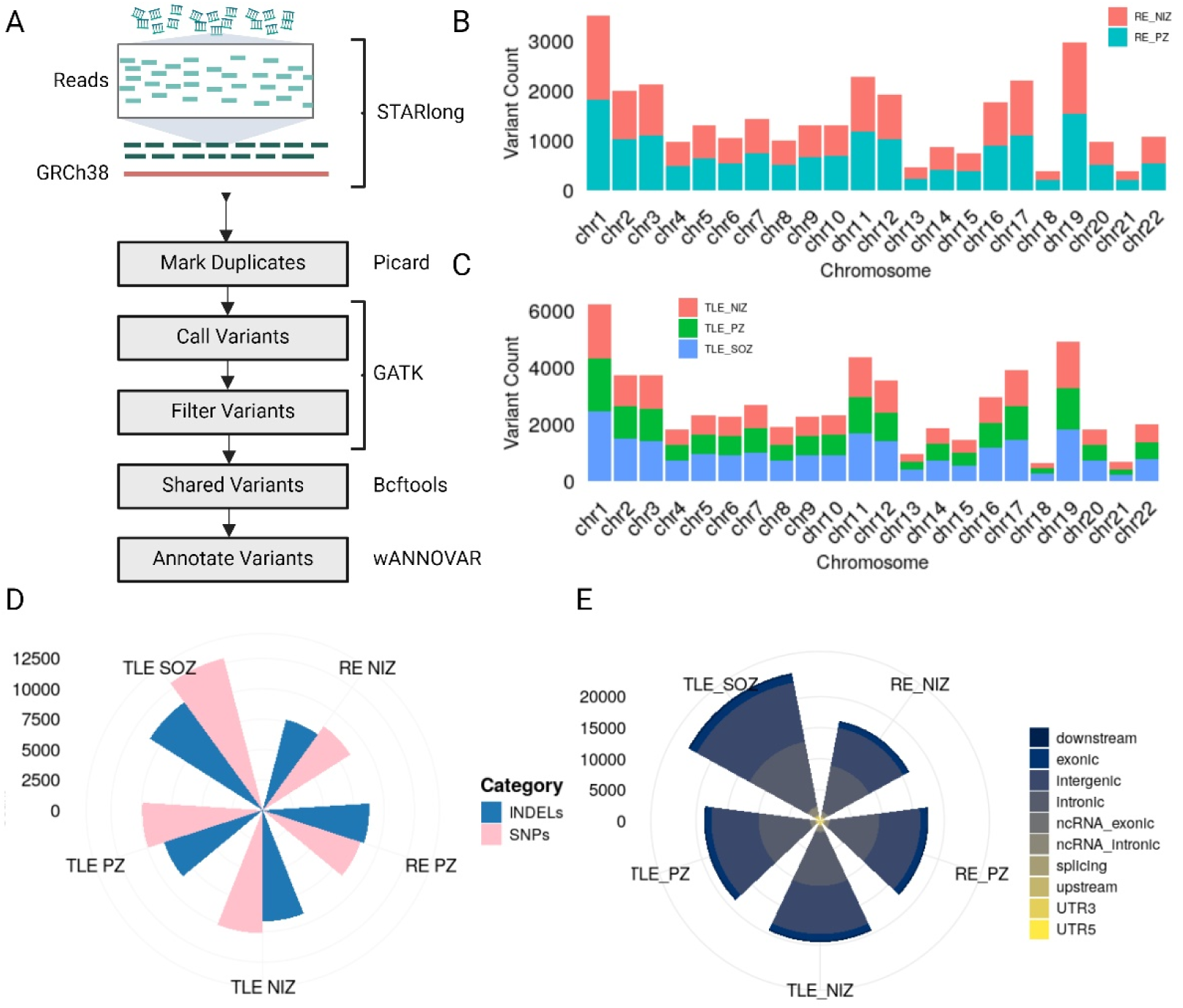
Identification of short variants in TLE and RE brains. (A) We identified short variants using GATK best practices workflows and annotated the high quality variants through wANNOVAR. (B-C). Chromosome-wise distribution of variants in RE (NIZ and PZ) and TLE (NIZ, PZ and SOZ) brains. (D) Number of variants falling in different categories in TLE and RE brains. (E) Functional annotations of variants harbouring at different genomic regions.

Using the transcriptomic data from the TLE samples, we identified a total of 4396 genes harbouring the variants in NIZ and among these 2.8% were unique to the NIZ region. In the SOZ region, we found a total of 4698 genes, among which were 9.5% specific to the SOZ region (Supplementary Figures 11A and B). In the PZ region, we identified a total of 4226 genes and among these 1.6% were unique to the PZ. A total of 3588 genes (59.7%) were shared across NIZ, PZ and SOZ (Supplementary Figures 11A and B). We then overlaid these variants with known epilepsy-associated genes^28^ and this analysis revealed that 3.9%, 3.9% and 4.6% of genes are shared among NIZ, PZ and SOZ respectively (Supplementary Figure S11A). Next, we performed GO-term analysis for the 3799 genes that were shared across the NIZ, PZ and SOZ. Interestingly this analysis showed that these novel variants were enriched for RISC complex assembly, mRNA, pre-RNA processing and RNA secondary structure unwinding pathways (Supplementary Figures 11C). NIZ-specific genes were enriched for negative regulation of inflammatory responses to antigens, negative regulation of DNA biosynthesis and regulation of intracellular signalling (Supplementary Figures 11D). For the PZ region, genes were enriched for the regulation of cellular localisation, apoptotic signalling pathways and protein deubiquitylation-related GO-terms (Supplementary Figures 11E). The NIZ genes were enriched for intracellular pH elevation, nuclear pore organisation and assembly and glial cell differentiation GO-terms (Supplementary Figures 11F). We then analysed the total number of genetic variants (INDELs and SNPs) present in SOZ, PZ and NIZ and found that SOZ harboured more genetic variants compared to PZ and NIZ (Fig 7D).

We also overlapped genes harbouring the variants with known epilepsy associated genes and found that 3.6% of genes were commonly present in PZ and NIZ (Supplementary Figures 12A). Similarly, in the RE brain, we found a total of 4095 genes harbouring the variants (517 unique) in the PZ area, and 4036 gene variants (458 unique) in the NIZ area among these variants 3578 are present in both NIZ and PZ (Supplementary Figures 11B). We further annotated these genes using GO enrichment analysis (Supplementary Figure 12C-E). In RE samples, the shared genes were enriched for protein localisation to chromatin, cytoskeletal organisation, and ribonucleotides biosynthesis (Supplementary Figure 12C), whereas the genes specific to the PZ regions were enriched for glycerophospholipid metabolism, lymphoid progenitor cell differentiation and axonal guidance-related processes (Supplementary Figure 12D). Furthermore, for the NIZ-related genes, mRNA processing and phosphorylation-related GO-terms were enriched (Supplementary Figure 12E). We then analysed the total number of genetic variants (Indels and SNPs) present in SOZ, PZ and NIZ and found the SOZ harboured more genetic variants compared to PZ and NIZ (Fig 7E). These observations suggest that SEEG electrode-derived RNA can detect genomic variants in addition to providing gene expression and DNA methylation profiles together with the matching electrophysiological and radiological data.

## DISCUSSION

Epilepsy is a complex neurological disorder characterized by recurrent seizures, with diverse aetiologies ranging from genetic mutations to environmental factors. Unravelling the molecular mechanisms underlying epilepsy etiology presents several challenges due to its multifactorial nature and heterogeneity. Genetic complexity, including polygenic inheritance and gene-environment interactions, complicates the identification of causal factors.^31–34^ Additionally, the dynamic nature of epileptogenesis and the interplay between excitatory and inhibitory neurotransmission further obscure our understanding.^35–37^ Limited access to human brain tissue, especially from living patients, hinders direct investigation of epileptogenic mechanisms. Furthermore, an incomplete understanding of seizure onset and network recruitment mechanisms and the lack of biomarkers for early diagnosis pose significant obstacles.^31, 33, 38, 39^ Addressing these challenges requires interdisciplinary collaboration, advanced experimental models and innovative technologies to decipher the intricate molecular landscape of epilepsy and develop targeted therapies.

Advances in neuroscience necessitate systems-level approaches that integrate multimodal data to understand the intricate structure and function of the human brain, particularly in health and disease.^40^ While studies utilizing postmortem tissue have provided valuable insights, accessing living brain tissue is essential for functional characterization. Treatment-resistant epilepsies offer a unique opportunity for such access, with *ex vivo* analysis of surgically-resected material enabling significant advances in understanding brain function.^41–43^ Intracranial EEG recordings from depth electrodes provides detailed spatial and temporal mapping of epileptogenic tissue, offering valuable insights into seizure onset and propagation. Recent studies have demonstrated the feasibility of extracting nucleic acids from depth electrodes, opening new avenues for molecular profiling of the human brain.^44, 45^

In this study, we demonstrate a novel method we call MoPEDE to extract nucleic acids from depth electrodes implanted in patients with epilepsy, enabling multimodal profiling of the transcriptome and epigenome in relation to the electrophysiological readings and radiological measurements. The utilization of SEEG electrodes, originally designed for electrophysiological measurements, as a source of RNA and genomic DNA (gDNA) represents a paradigm shift in epilepsy research and the broader field of network neuroscience. Through meticulous experimental procedures, including electrode sectioning, nucleic acid extraction, and comprehensive profiling, we demonstrate the feasibility of obtaining high-quality RNA and gDNA from SEEG electrodes, even from deep brain structures. Our methods revealed distinct molecular landscapes associated with different epilepsy subtypes, including FCD, non-lesional TLE, and RE. The inclusion of these etiologies demonstrates MoPEDE applies across spatial scales ranging from highly localised (FCD) through to whole hemisphere (RE). We identified transcripts and DNA methylation profiles that correlate with recorded neurophysiological signals, highlighting the relevance of molecular profiling in understanding epilepsy aetiology and guiding treatment strategies. While the SEEG electrodes covered only selected regions of the brain, surgical outcomes are the gold standard for measuring successful identification and surgical removal of epileptogenic tissue. The excellent surgical outcome, along with accurate ictal recordings support that the electrodes were in the epileptic foci (Table 1). Additionally, the identification of specific genes and pathways enriched in epileptic brain regions, validated through comparison with publicly available datasets and single-cell RNA sequencing data,^46, 47^ reinforces the robustness and significance of the findings. Additionally, our approaches further demonstrate the ability to derive information about clinical variants, such as SNPs and Indels, from a limited number of SEEG electrodes. This includes identifying both pathogenic ^48^ and risk factor variants across various brain regions affected by epilepsy and their associated functions ^21^.

The significance of the variation in numbers of differentially expressed genes in relation to epileptogenicity index and etiologies is not fully understood but indicates that the trace nucleic acids on SEEG contacts reflect biological processes that differ in the SOZ and PZ compared to NIZ. This included a number of genes associated with cell metabolism, processes increasingly recognised across epilepsy etiologies. In both TLE and RE samples, we found differences in the SOZ compared to PZ that included transcripts regulating cell death. This could reflect the engagement of pathways that limit neurodegeneration in tissue actively generating seizures, processes known to be evoked by repeated seizures^49^. Analysis of SNPs and indels within the transcriptome data showed enrichment of many of the same pathways, as well as highlighting RNA processing and chromatin processes that are also implicated in the pathogenesis of epilepsy^50–53^. The amount of these variants appears to scale with EI score implying a causal effect of tissue pathogenicity as a driver as well as potential diagnostic applications in distinguishing SOZ from PZ. Further, DNA methylation findings showed alignment with findings from tissue-based studies of FCD and TLE, in terms of numbers of hyper- and hypo-methylated genes^54, 55^ and biological processes influenced by this epigenetic mark.

Correlations between SOZ and PZ suggest a proportion of epigenetic marks reflect the effects of seizures per se whereas others may distinguish seizure-triggering sites from recruited networks. Thus, the surface of explanted SEEG contacts bears molecular traces reflective of cellular biology and pathology of the source tissue.

The source of the detected transcripts and epigenetic signals on the electrode surface is likely to be from the surrounding neurons, glia and perhaps vascular cells at that specific site or along the path of contact. Indeed, we detected transcripts representing multiple resident brain cell types in all three cases. The cause of the release of these nucleic acids is likely mechanical injury due to insertion and/or withdrawal of the electrode, but controlled release is also possible. Indeed, both neurons and glia are capable of releasing packets of membrane-enclosed cellular material in the form of extracellular vesicles such as exosomes ^56^. While the extent and functional significance of such information-carrying paracrine signalling remains incompletely understood, experimentally evoked seizures have been reported to adjust the abundance and nucleic acid content of extracellular vesicles^57^. The coherence between epigenetic and transcript signals suggests these materials had a similar cellular source, but it is possible that origin and release mechanisms differ between the two nucleic acid types. Moreover, the source of some nucleic acids may be from infiltrating immune cells, which deliver such material to resident brain cells, including neurons after seizures^58^. While RE is most strongly associated with an infiltrating inflammatory cellular presence, both TLE,^59^ and FCD^60^, feature immune cell infiltration. Further studies may yield a more complete understanding of the basis of the detected signals and their relationship to pathophysiological communication from local and perhaps non-resident cellular sources.

There are a number of limitations to consider in the present study. Obtaining sufficient quantities of high-quality nucleic acids from a limited number of SEEG electrodes remains challenging for NGS methods. This may be addressed by further improvements in the coordination of workflow for on-site sample collection. For example, reducing the time to snap-freezing electrodes after their removal or proceeding to immediate extraction of nucleic acids may enhance both the quality and quantity of retrieved material. Indeed, with the current extraction method, we did not achieve uniform sample integrity with several samples showing degradation and failing to pass the QC in the NGS workflow. Additional or new extraction methods should better ensure the preservation of sample integrity. Second, other information-containing material may also be recoverable. For example, histone modifications which play a major role in controlling gene expression and cellular identity^38, 61^. Profiling histone modification changes between different epilepsy subtypes may complement MoPEDE and extend insights into disease pathophysiology and etiology. Extracting sufficient histones from SEEG electrodes may require the development of new methods.

Last, SEEG has an inherent sampling bias with low spatial resolution, being able to cover only a limited area of the brain or epileptic network. Increasing numbers of depth electrodes must be balanced with limiting intra-operative and post-operative complications of multiple-depth electrode placement. Visual inspection of SEEG data is open to reader variability, and while still the clinical standard, has prompted the development of more automated techniques such as the EI. EI scores are more accurate with fast frequencies at seizure onset but are not as accurate with slower frequencies, for example, the temporal pole SOZ in patient B had a slow ictal onset frequency with a low EI score. EI does not uniformly correlate with visual inspection. For example, EI scores in the RE case were diffusely elevated, including in the NIZ, likely due to the expected diffuse epileptogenicity in the affected hemisphere.

In conclusion, our findings offer promising prospects for improving the diagnosis and treatment of epilepsy. By integrating EEG-based epileptogenicity index scores with multiomic data, we gain deeper insights into disease mechanisms, paving the way for more targeted and effective therapies. Furthermore, the ability to extract nucleic acids from depth electrodes provides a non-invasive means of molecular profiling, offering potential applications beyond epilepsy, such as studying other neurological disorders. Future research directions may involve exploring additional nucleic acids (e.g. long non-coding RNA, small RNAs), epi-transcriptome changes (RNA modifications) and epigenetic marks (histone marks) obtained from depth electrodes, as well as investigating the applicability of this approach across diverse epilepsy subtypes.

## MATERIALS AND METHODS

### Ethics Statement

The present study was reviewed and approved by the Beaumont Hospital Medical Research Ethics Committee under study no. 20.58. All patients provided informed consent.

### Neurophysiology data collection

All participants underwent robot-assisted implantation of intracranial depth electrodes and SEEG monitoring in the Epilepsy Monitoring Unit in Beaumont Hospital Dublin, Ireland. Depth electrodes with either twelve, fifteen, or eighteen contacts (DIXI Medical, France) were implanted according to the pre-operative hypothesis of the seizure-onset zone. By standard nomenclature, each depth is labelled with a letter, with electrode contacts numbered from mesial to lateral. The anatomic location of each electrode was confirmed by postoperative CT co-registered with preoperative volumetric MRI (Figure 2). Continuous EEG was recorded at 1024Hz with concurrent video recording (Xltek EEG System, Natus Inc.). Two experienced clinical neurophysiologists/epileptologists performed visual inspection (VI) of seizures and classified each cortical region as part of the SOZ, PZ, or NIZ, , based on summation of ictal EEG data. The SOZ is defined as the electrodes involved at the onset of the EEG seizure, and correlates with an epileptogenicity index (EI) >0.4. The PZ is defined as the electrodes involved in early seizure propagation within 10 seconds, and correlates with an EI of 0.2-0.4. The NIZ is defined as electrodes not involved in the seizure and correlates with an EI<0.2.

To measure the epileptogenicity index (EI), EEG seizures were analysed using the AnyWave software (Marseille, France).^24, 25^ Representative seizures from each patient which occurred over forty-eight hours from electrode implantation and with minimal artefact were selected. Bipolar contacts in grey matter at the mesial or lateral point of each depth electrode represented each brain region under investigation. Contacts in white matter or outside of the brain were excluded. The EI measures the change in energy ratio and time delay from electrode contacts at seizure onset to estimate the epileptogenicity of a given cortical region.

The depth electrodes were explanted under general anaesthesia following standard clinical procedures. They were immediately placed in sterile, RNA, DNA and nuclease-free 15ml tubes and frozen in dry ice for transport. The electrodes were then stored at -80°C until further processing.

### Patient clinical characteristics

Patient A is a 49-year-old male who underwent SEEG monitoring for medication-resistant focal epilepsy due to a right parietal focal cortical dysplasia (FCD). Thirty-three electroclinical seizures were recorded. The EEG onset for all seizures arose from the lateral contacts of the X electrode, at the parietal FCD. Seizure propagation involved the supplementary motor area, and temporo-occipital junction. The patient underwent resection of the FCD and achieved an Engel class 1A outcome, remaining seizure-free at one year. Histopathology confirmed a type IIa FCD.

Patient B is a 40-year-old female with MRI-negative, medication-resistant temporal lobe epilepsy who underwent SEEG monitoring. Six electroclinical seizures were recorded. EEG onset was from the anterior hippocampus, and the temporal pole. Seizure propagation involved the middle temporal gyrus and inferior temporal gyrus. The patient underwent a left temporal lobectomy with amygdalo-hippocampectomy, resulting in an Engel class 2A outcome at one year, initially seizure-free but experiencing rare seizures subsequently. Histopathology revealed Chaslin’s subpial gliosis.

Patient C is a 24-year-old female with Rasmussen’s encephalitis at the age of 15 years treated medically. Due to persistent medication resistant epilepsy, she underwent SEEG monitoring. Thirty-two electroclinical seizures were recorded. Ictal onset of most seizures was from the hippocampus and the gyrus rectus. Seizure propagation involved the cingulate, and frontal operculum. The patient underwent right frontal and anterior temporal resection with an Engel class 2A outcome at one year, initially seizure-free with rare seizures subsequently. Initial brain biopsy at age 15 years showed chronic encephalitis with T-cell rich perivascular and parenchymal inflammation. Histopathology from the resection eight years later revealed severe hippocampal sclerosis and cortical gliosis, with no active inflammation. (Table 1)

### Total Nucleic acid extraction

For the total nucleic acid extraction, snap-frozen SEEG electrodes were cut into small pieces using sterile nuclease-free scissors into microfuge tubes and the total nucleic acids were extracted using PicoPure™ RNA Isolation Kit (Thermo KIT0204) according to the manufacturer’s instructions with minor modifications. Briefly, 50 to 100µl (up to the electrode pieces were completely immersed) extraction buffer was added to the cut electrodes followed by 30 minutes of incubation at 42°C. Meanwhile, the column was preconditioned with 250µl of Conditioning Buffer. An equal amount (50 to 100µl) of 70% ethanol was added to the extracted samples and mixed, transferred to the preconditioned column centrifuge for 2 minutes at 100 x g, immediately followed by centrifugation at 16,000 x g for 30 seconds to remove flowthrough. Bound fractions were washed with wash buffer I without DNase to retain the genomic DNA in the same samples followed by wash buffer II and the samples were eluted with 11µl of elution buffer.

Samples were quantified in a nanodrop and the quality of the samples was analysed using an Agilent TapeStation system with high-sensitivity RNA tapes or in a fragment analyser. For the separation of DNA and RNA, the eluted samples were divided into equal portions and one portion was used for RNA isolation and the other was used for DNA isolation.

### RNA purification

For the RNA purification samples were first treated with DNase I (Thermo EN0521) to remove the DNA and purified using GeneJET RNA Purification Kit (Thermo K0732) according to the manufacturer’s instructions. Purified RNA samples were quantified using Qubit high-sensitivity RNA Quantification assay (Thermo Q32852). The quality of the RNA was analysed using Agilent TapeStation systems with high-sensitivity RNA tapes with RIN numbers.

### Genomic DNA purification

For gDNA purification samples were added to nuclease-free water (Thermo AM9939) to the final volume of 100µl. Then 3µl of RNase A (Thermo R1253) was added to the samples and incubated for 20 minutes at 37°C. Further DNA was purified using Monarch® Genomic DNA Purification Kit (NEB T3010S) according to the manufacturer’s instructions with minor modifications. 1µl of Proteinase K (Thermo EO0491) was added to the samples and 100µl of lysis buffer was added and incubated for 5 min at 57°C followed by 400µl of binding buffer was added and loaded to the purification columns. Washed twice with wash buffer and eluted with 15µl of elution buffer. Purified DNA samples were quantified using Qubit high-sensitivity dsDNA Quantification assay (Thermo Q32854). The quality of the gDNA was analysed using an Agilent TapeStation system with high-sensitivity genomic DNA tapes.

### Bulk FLASH-seq

Following extraction, RNA samples were normalized to 1ng/uL and 1 μL of RNA was input to prepare RNA libraries using a bulk input optimized FLASH-seq protocol.^27^ In brief, RNA was converted to cDNA fragments using Maxima H Minus (#EP0753, Thermo Fisher Scientific) and amplified with KAPA HiFi HotStart (#KK2602, Roche). Note, double lysis mix volume used and 4uL RTPCR mix used to account for larger wells in 96 well plate. cDNA was then cleaned using a 0.8x ratio of homebrew SeraMag beads in 18% PEG (#GE24152105050250, CytiviaTm). cDNA concentrations and sizings were checked using Qubit (#Q33231, Thermo Fisher Scientific) and Agilent Bioanalyzer (#5067-4626 Agilent). cDNA were normalized to 200 pg/μL before tagmentation using 0.2 μM of homemade Tn5 (EPFL). The reaction was halted with 0.2% SDS. Indexing PCR was performed to add Nextera index adapters (1 μM final, Integrated DNA Technology) using KAPA HiFi reagents (#KK2102, Roche). Libraries were pooled in equal volumes and a final 0.8x cleanup was performed with homebrew SeraMag beads before measuring the sample concentration and sizing. The library pool was normalized and sequenced on Illumina NovaSeq SP flowcell (75-8-8-75) at approximately 25 million reads/sample. Basecalling and demultiplexing were performed with bcl2fastq (v2.20, Illumina Inc.).

### Transcriptome quality check, mapping, quantification and variant analysis

Adapter sequences were removed from raw FASTQ files using Cutadapt (v4.9).^62^ Subsequently, high-quality, adapter-trimmed reads were mapped against the human reference genome (GRCh38) using the STARlong utility within the STAR aligner.^63^ Transcriptome assembly and quantification of gene expression was performed using StringTie2 at default parameters for each electrode from different brain regions in each case.^64^ Then we performed differential gene expression for all regions and epilepsy brains using DESeq2.^65^ We incorporated publicly available single-cell RNA-seq (scRNA-seq) data from epilepsy patients,^23^ into our analysis. Quality checks, data reduction, and integration of healthy and epilepsy scRNA-seq data were performed using Seurat (v4).^47, 66^ To identify variants using transcriptome data, we employed the GATK (v4.5.0.0) RNAseq short variant discovery (SNPs + Indels) pipeline.^47^ The process began by marking duplicate reads with MarkDuplicates. Next, we used SplitNCigarReads to split reads with N in the CIGAR string into multiple supplementary alignments and hard clip mismatching overhangs. We then used AddOrReplaceReadGroups (Picard) to assign all reads in a file to a single new read group. Variants were called using HaplotypeCaller, followed by high-quality variant filtering with VariantFiltration, applying filters at QD < 2.0, FS > 60.0, MQ < 30.0, DP < 20.0, and QUAL < 20.0.^46^ Variants effect was determined using Ensembl Variant Effect Predictor (VEP) for high quality variants.^67^ Further we intersect the variants present in all replicate for a region in epilepsy brains using BCFtools,^68^ and annotated them using wANNOVAR.^69^

### Methylation quality check, mapping, and differential Analysis

We performed quality checks using FastQC to assess the overall quality of the bisulfite-converted sequencing (BS-Seq) reads. Following quality control, adapter sequences were trimmed using TrimGalore. The human reference genome (GRCh38) was prepared using the bismark_genome_preparation utility from Bismark to account for bisulfite conversion. Subsequently, high-quality, adapter-trimmed reads were mapped to the prepared reference genome using Bismark (v0.22.3).^70^ Methylated cytosines were extracted from the mapped reads using MethylDackel, considering all three methylation contexts: CpG, CHG, and CHH (dpryan79/MethylDackel: A (mostly) universal methylation extractor for BS-seq experiments GitHub repository). Differentially methylated regions (DMRs) were identified using methylKit with a sliding window approach (window size = 200 bp, step size = 200 bp, minimum coverage = 10 bp, difference ≥ 5 & ≤ -5 and q value ≤ 0.05 ) using default parameters,^71^ Methylated regions were further annotated using the bedtools intersect utility,^72^ to identify overlapping genes. Finally, we integrated methylation information at the gene level with their corresponding transcriptional expression data.

### Visualization and statistical analysis

We used mapped BAM files to generate BigWig files for visualization of transcriptome and methylome density in gene bodies and flanking regions using deepTools,^73^ various utilities available within SAMtools to prepare the file inputs for different analyses.^74^ Statistical analyses and visualizations were performed using R packages (R Core Team, 2023). Finally, BioRender was used to create schematics and enhance the visual presentation of our figures.

## AUTHOR CONTRIBUTIONS

AD performed data analysis and wrote the manuscript. AM performed molecular analysis of samples and wrote the manuscript; AS and JL participated in sample procurement, neurophysiology and clinical data collection and analysis. KJS and DFO’B performed SEEG implantation procedure, surgical resections and obtained consent. PWW oversaw and interpreted the clinical, neurophysiological, radiological and pathological data collection and analysis, and edited the manuscript. RS and SP performed the FLASH-seq procedure on SEEG-derived RNA. VKT and DCH co-conceived the study, obtained funding, supervised the study, performed data interpretation, and co-edited the manuscript.

## ACKNOWLEDGEMENTS

The authors thank the patients for the donation of samples for the present study. We thank the iCLOUD team at SDU and HPC team at QUB for managing the HPC cluster and configuring storage, cluster job-queues, installing software and giving helpful advice. We thank the QUB Genomics facility for assistance with DNA Methylome data generation. The authors thank Karina Halley for support with ethics and Austin Lacey for technical support with sample collection. We also thank Ronan Kilbride, Department of Clinical Neurophysiology, Beaumont Hospital, Dublin for additional visual inspection of SEEG data and Alan Beausang, Department of Neuropathology for the provision and interpretation of neuropathology slides and our clinical colleagues at Beaumont Hospital. This publication has emanated from research supported by a research grant from the Higher Education Authority Ireland (SeeDeepER) jointly to DCH and VKT, Science Foundation Ireland (SFI) under grant 16/RC/3948 and 21/RC/10294_P2 (FutureNeuro) and European Union Horizon 2020 (FET-Open award 964712) to DCH, and Novo Nordisk Foundation 3110103 and Danish National Research Foundation DNRF177 grants to VKT.

## CONFLICT OF INTEREST

PWW has been a paid speaker for Jazz Pharmaceuticals (epidiolex) and Angelini (ontozry). Remaining authors declare no conflicts of interest.

## Data availability

We have submitted raw fastq and count matrix for transcriptome and methylome data generated in this study to Gene Expression Omnibus (GEO) using accession numbers GSE268714 and GSE268715.

---

**Figure S1.**
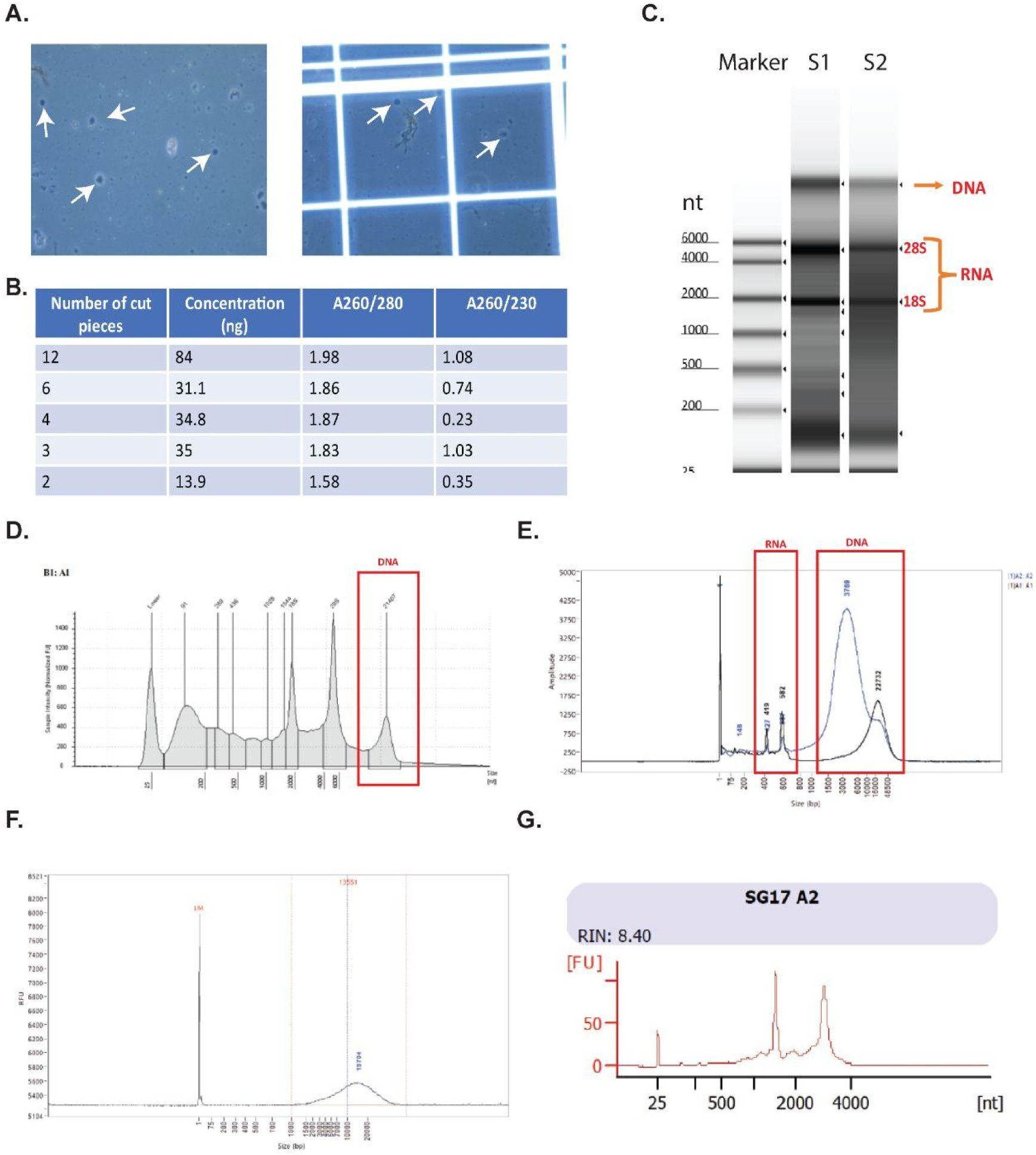
Nucleic acid extraction from SEEG electrodes from epilepsy patients. (A). Trypan blue staining shows the presence of cells in the SEEG electrodes from epilepsy patients. (B). The amount of total nuclear fractions was directly proportional to a number of cut pieces (Metal contact points) from one electrode. (C). High sensitivity fragment analyser shows the presence of both DNA and RNA in the samples (S1 and S2). (D-E). Electropherograms from the fragment analyzer confirm the RNA and DNA with their respective molecular weight. (F). Electropherograms from the fragment analyzer confirm the effective separation of DNA from total nucleic acid fraction. (G). Tape station analysis of purified RNA fraction confirmed the quality of (RIN) extracted RNA.

**Figure S2.**
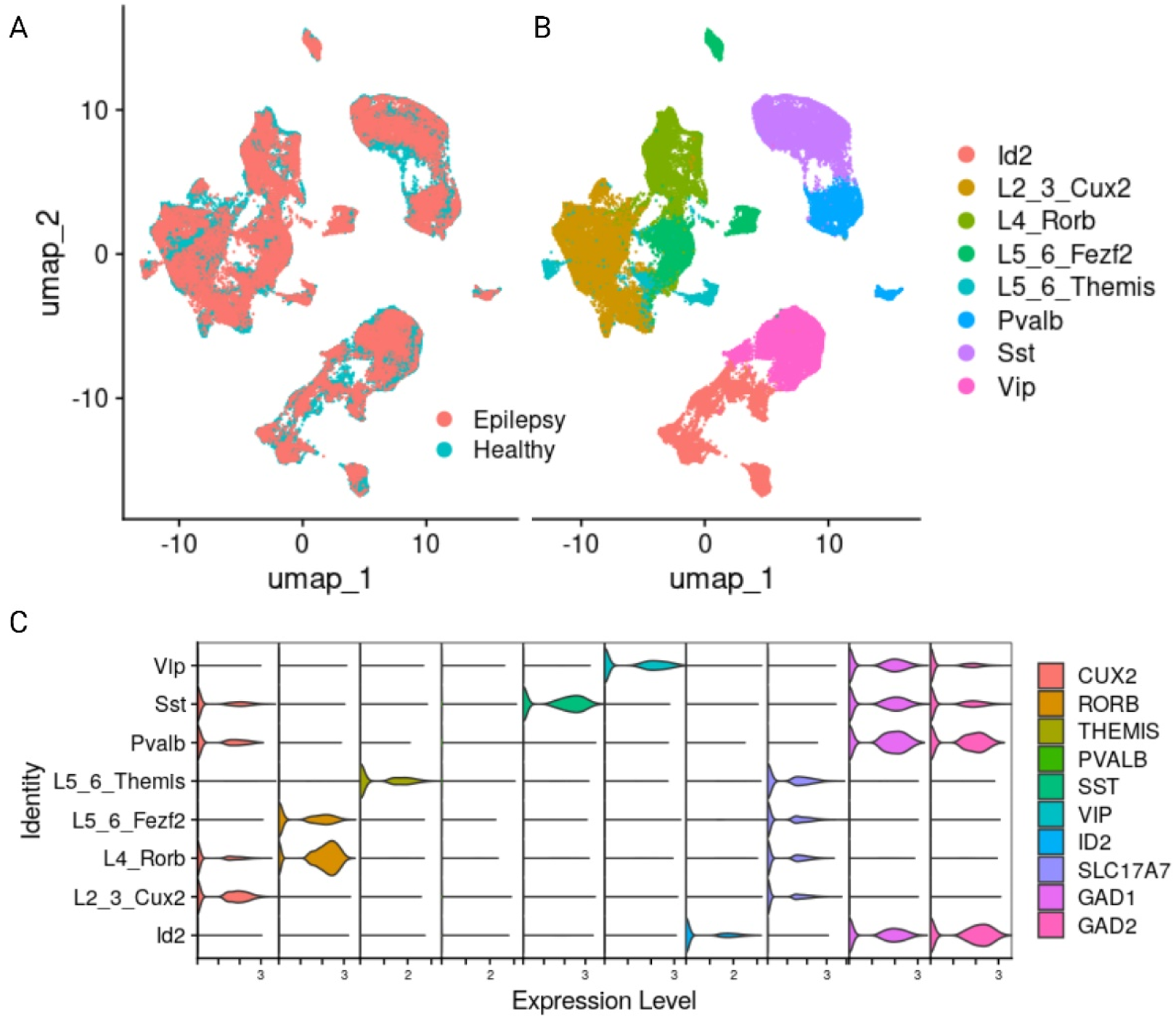
Epilepsy and control brain scRNA-seq data used for comparative analysis. (A) UMAP shows the integration of epilepsy and healthy brain scRNA and (B) their cell type annotation. (C) Expression of markers gene to different cell types.

**Figure S3.**
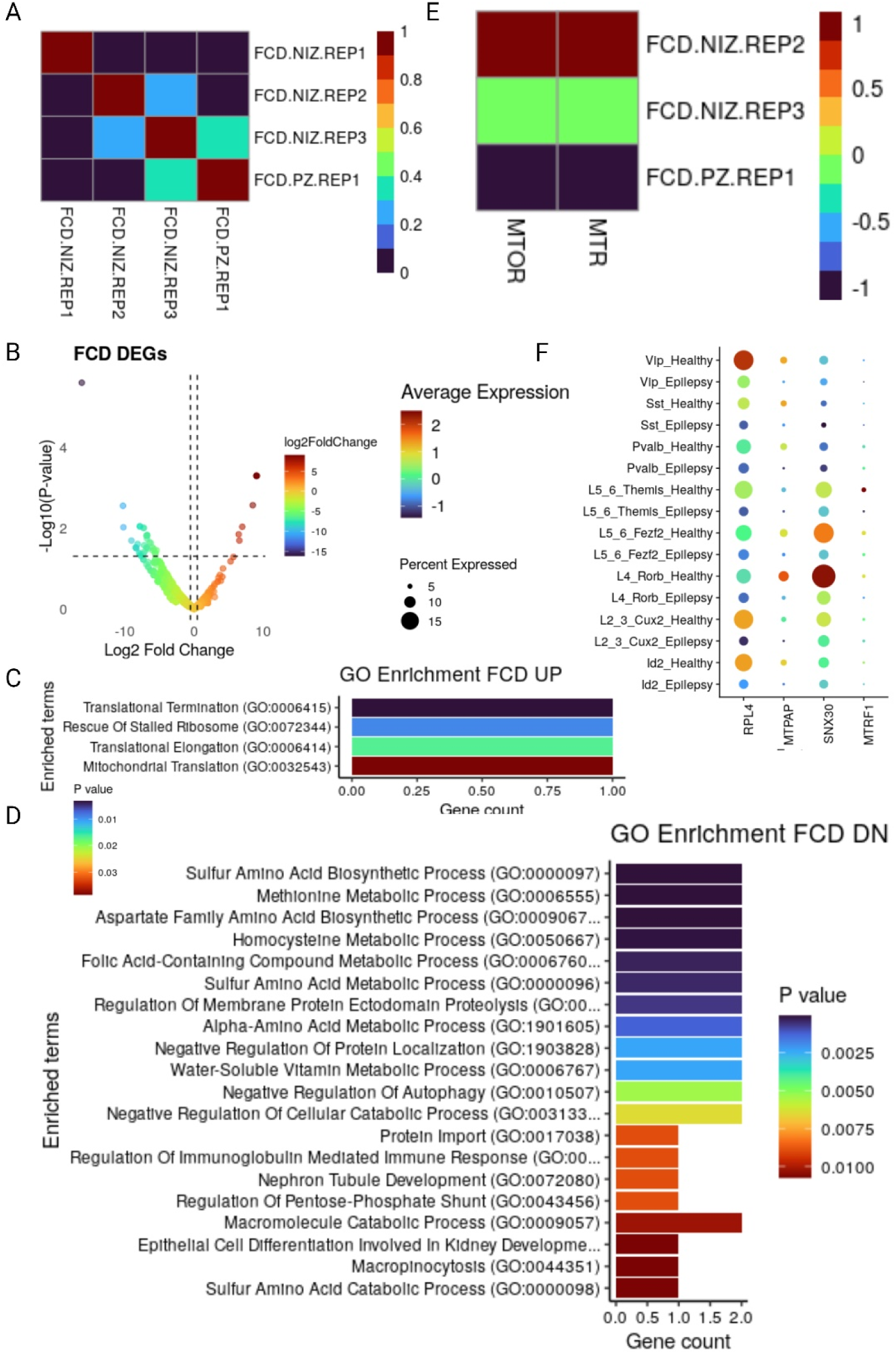
FCD transcriptome response in NIZ and PZ areas. (A) Replicate-wise correlation of NIZ and PZ regions transcriptome. (B) Volcano plot of differentially expressed genes between PZ and NIZ regions. (C-D) GO enrichment of up and down regulated genes, respectively. (E) Heatmap of down regulated known epilepsy risk genes in PZ as compared with NIZ region. (F) Dotplot showing the expression of down regulated genes in PZ vs NIZ and Epilepsy vs Healthy from public data.

**Figure S4.**
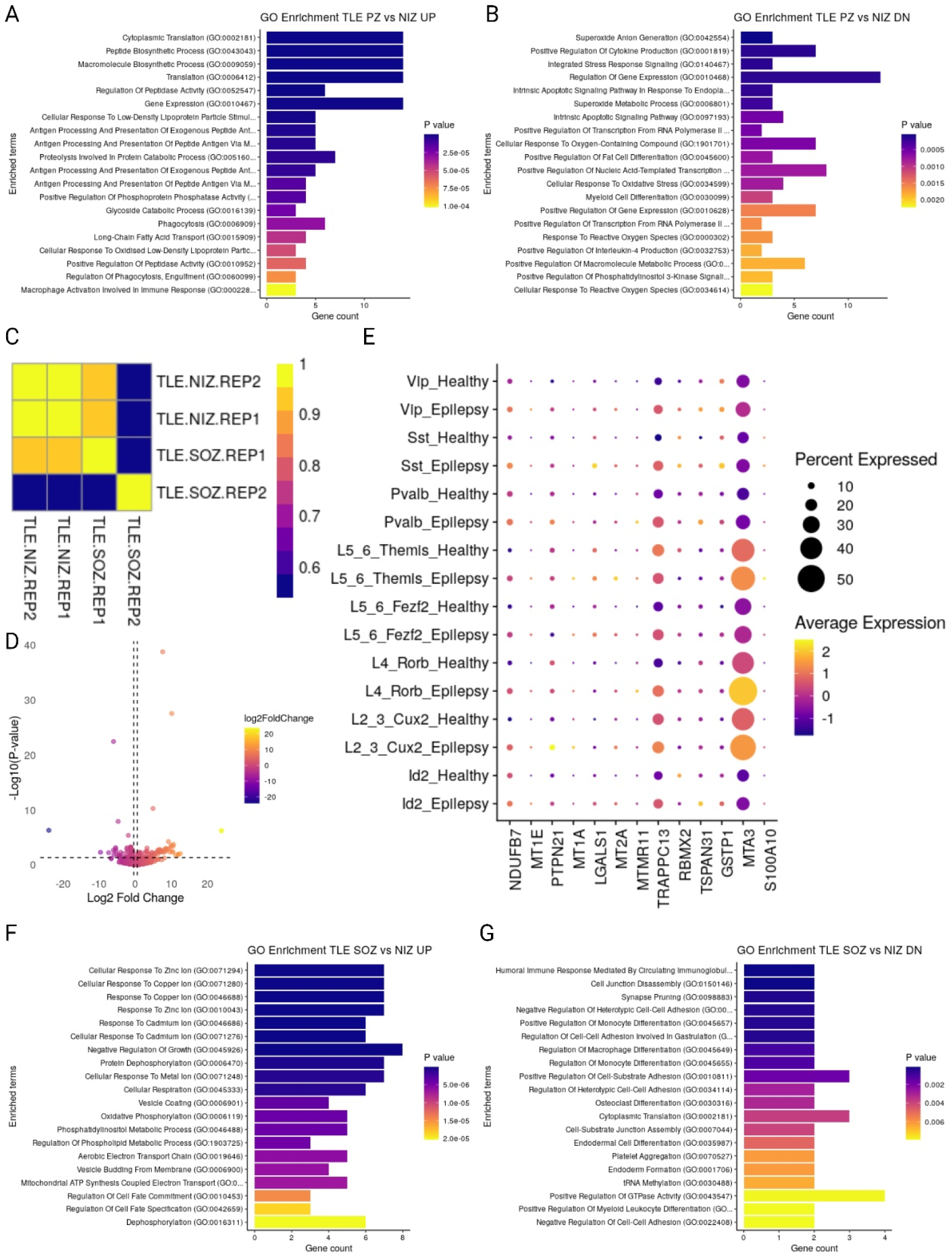
TLE transcriptome response in NIZ and SOZ areas. This figure examines gene expression changes in different regions of non-lesional temporal lobe epilepsy (TLE) brains. Panel (A-B) GO enrichment for up- and downregulated genes in PZ vs NIZ comparison in TLE brain, respectively. (C-D) Replicate wise correlation of NIZ and SOZ regions and volcano plot of differentially expressed genes for the same regions. (F-G) GO enrichment for up- and downregulated genes in SOZ vs NIZ comparison in TLE brain, respectively. (E) Dotplot showing the expression of upregulated genes in SOZ vs NIZ and Epilepsy vs Healthy from public data.

**Figure S5.**
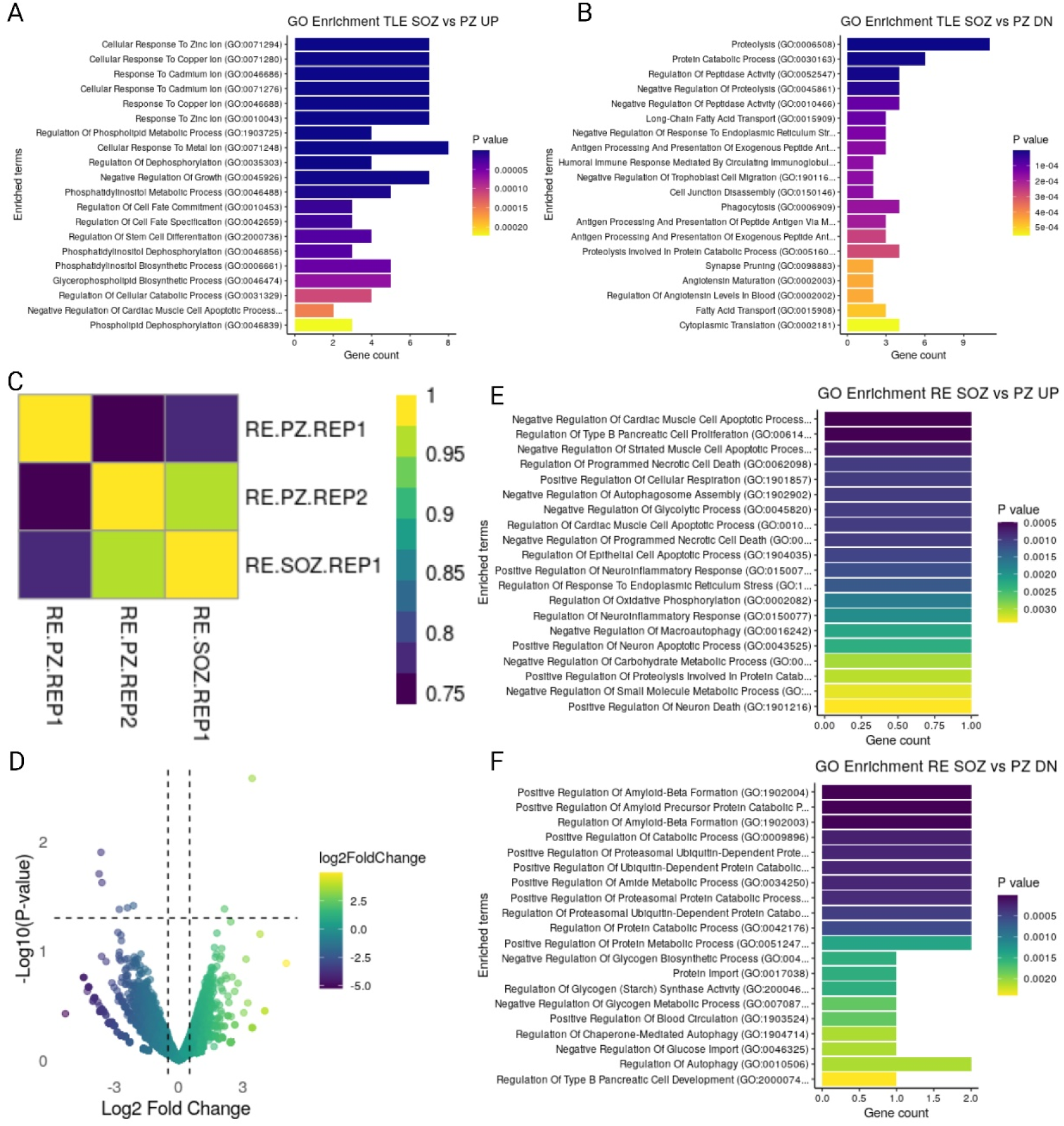
Transcriptome response in PZ and SOZ areas. This figure examines gene expression changes in PZ and SOZ of TLE and RE brains. Panel (A-B) GO enrichment of up- and downregulated genes in SOZ vs NIZ comparison in TLE brain. (C-D) Replicate-wise correlations and differential expression of genes in SOZ and PZ regions of RE brain. (E-F) GO enrichment of up- and downregulated genes in SOZ vs PZ region comparison of RE brain.

**Figure S6.**
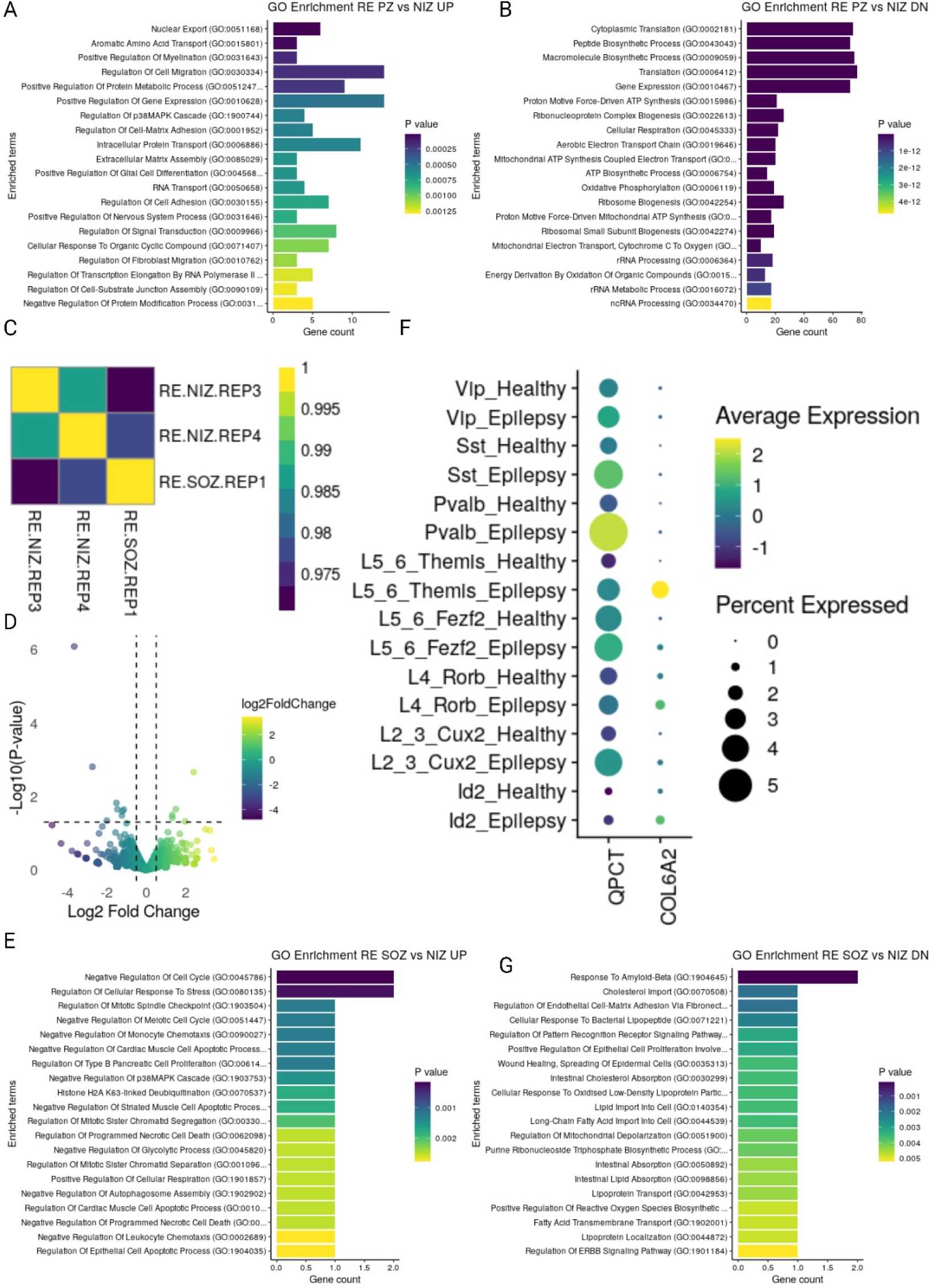
RE transcriptome response in NIZ, PZ and SOZ regions. (A-B) GO enrichment of up- and downregulated genes in PZ vs NIZ region comparison of RE brain. (C-D) Replicate-wise correlations and differential expression of genes in SOZ and NIZ regions of RE brain. (E and G) GO enrichment of up- and downregulated genes in SOZ vs PZ region comparison of RE brain. (F) Dotplot showing the expression of upregulated genes in SOZ vs NIZ and Epilepsy vs Healthy from public data.

**Figure S7.**
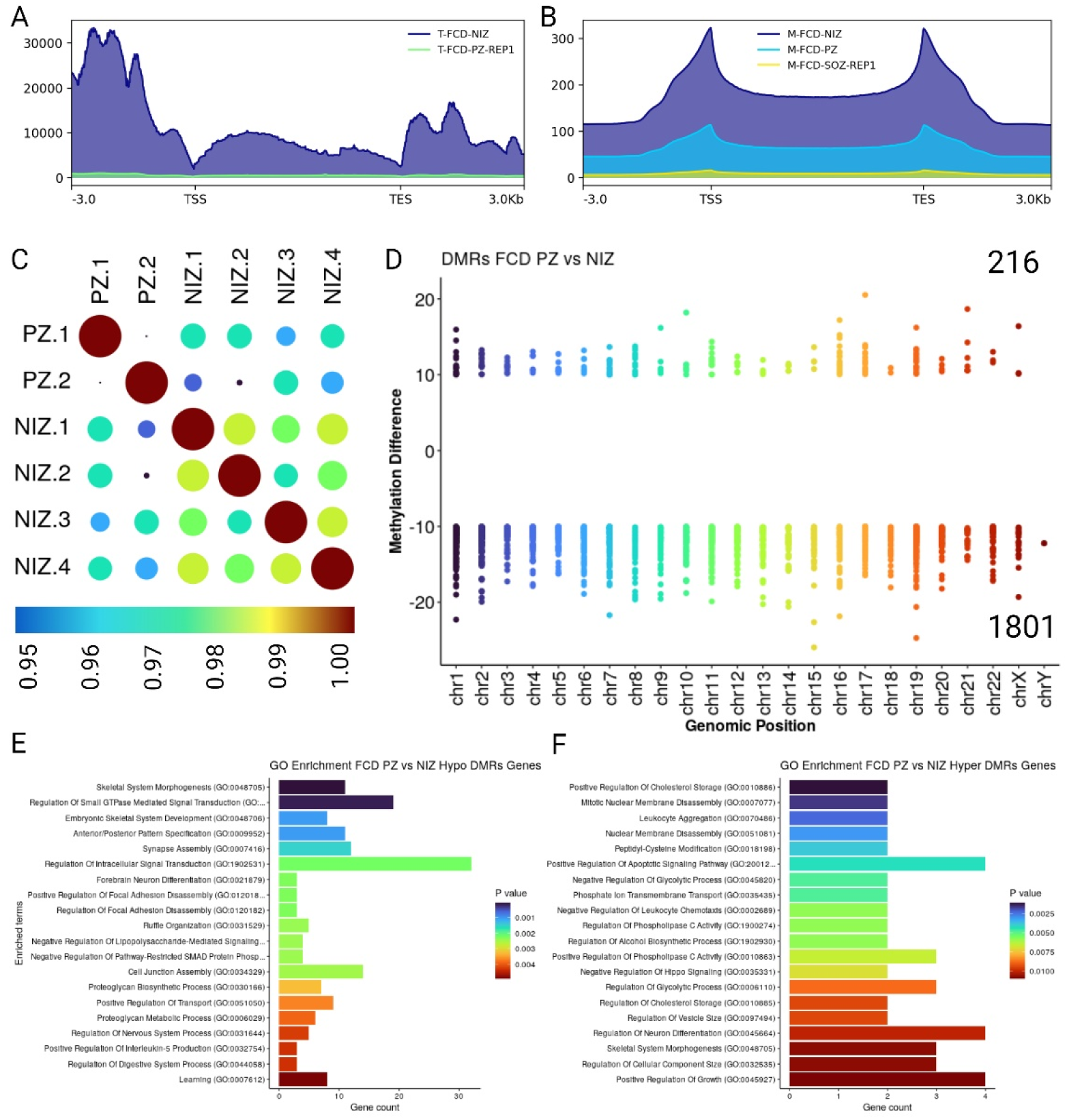
Methylome landscape in FCD. (A-B) Transcriptome and methylome density plot in gene body and flanking regions. (C) Replicated wise correlation plot of PZ and NIZ regions methylome in FCD brain. (D) A comparable number of hypermethylated (216) and hypomethylated (1801) DMRs are identified when comparing the PZ to the NIZ region. (E-F) GO enrichment of hypomethylated and hypermethylated genes found in PZ regions.

**Figure S8.**
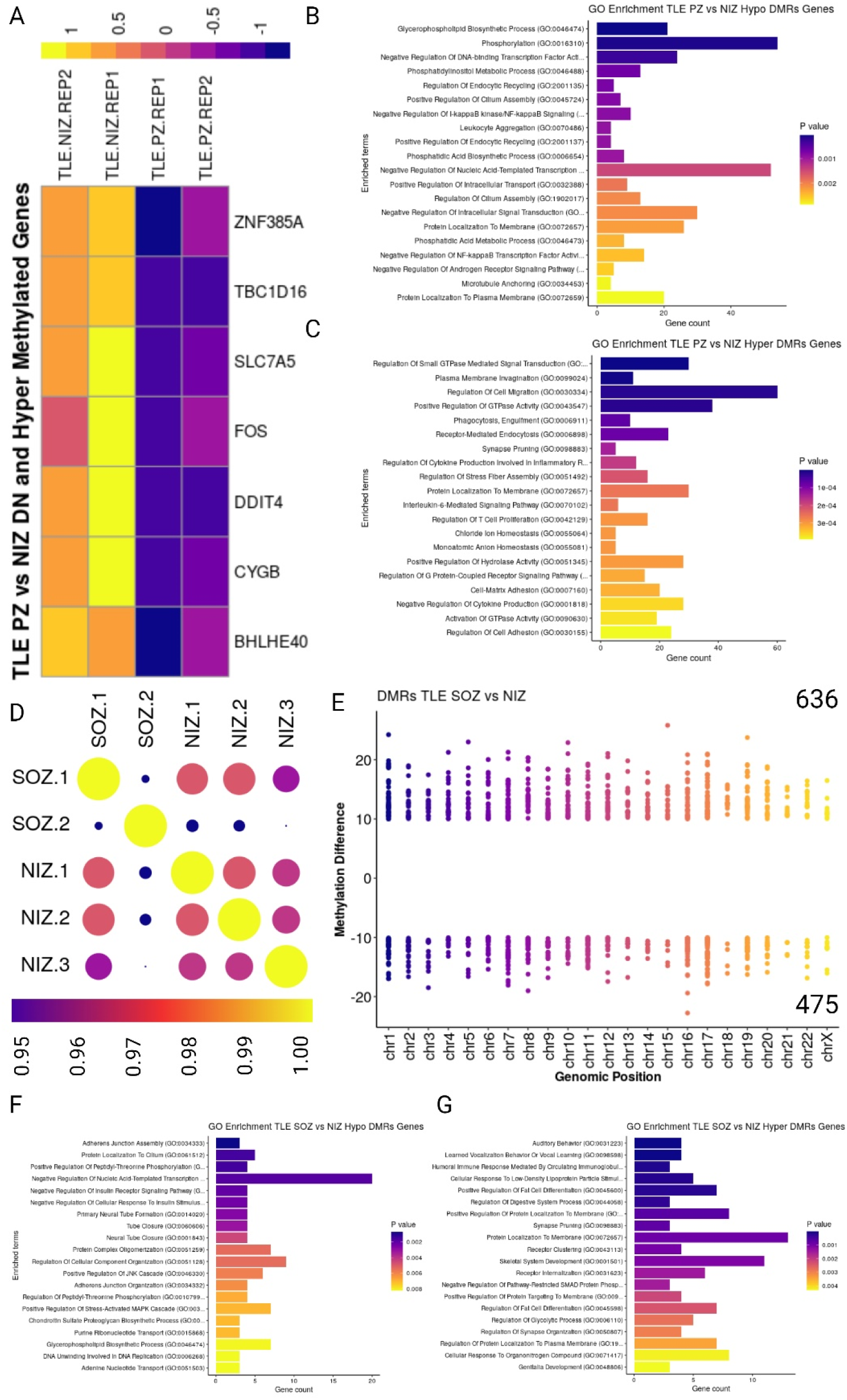
Transcriptome and methylome comparison in TLE. (A) Heatmap of down regulated and hypo methylated genes in PZ vs NIZ comparison. (B-C) GO enrichment of hypomethylated and hypermethylated genes found in PZ regions. (D) Replicate-wise correlation plot of SOZ and NIZ regions methylome in TLE brain. (E) A comparable number of hypermethylated (636) and hypomethylated (475) DMRs are identified when comparing the SOZ to the NIZ region. (F-G) GO enrichment of hypomethylated and hypermethylated genes found in SOZ regions.

**Figure S9.**
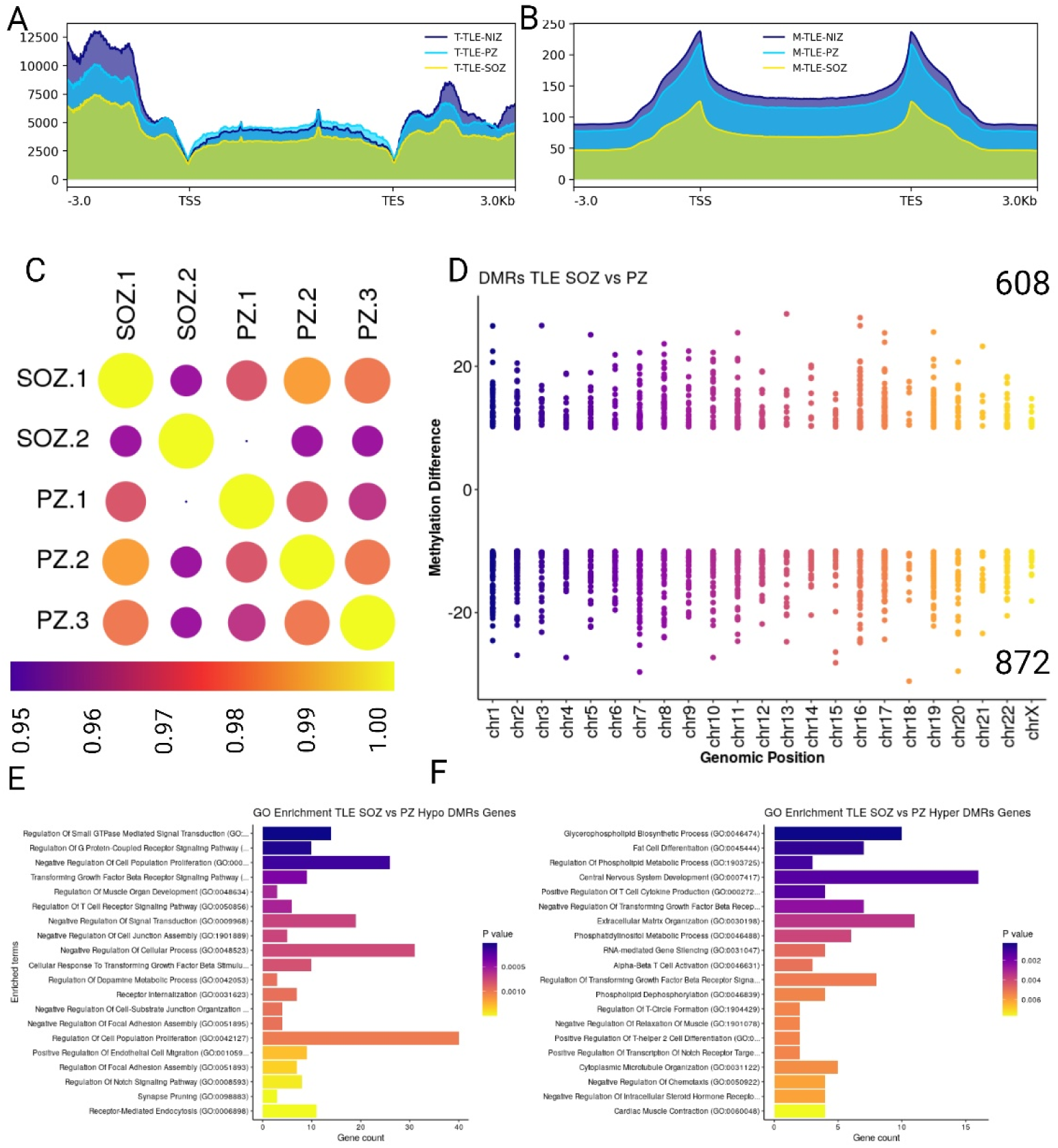
Transcriptome and methylome analogy in SOZ and PZ regions in TLE. (A-B) Transcriptome and methylome density plot in gene body and flanking regions. (C) Replicated wise correlation plot of SOZ and PZ regions methylome in TLE brain. (D) A comparable number of hypermethylated (608) and hypomethylated (872) DMRs are identified when comparing the SOZ to the PZ region. (E-F) GO enrichment of hypomethylated and hypermethylated genes found in SOZ regions.

**Figure S10.**
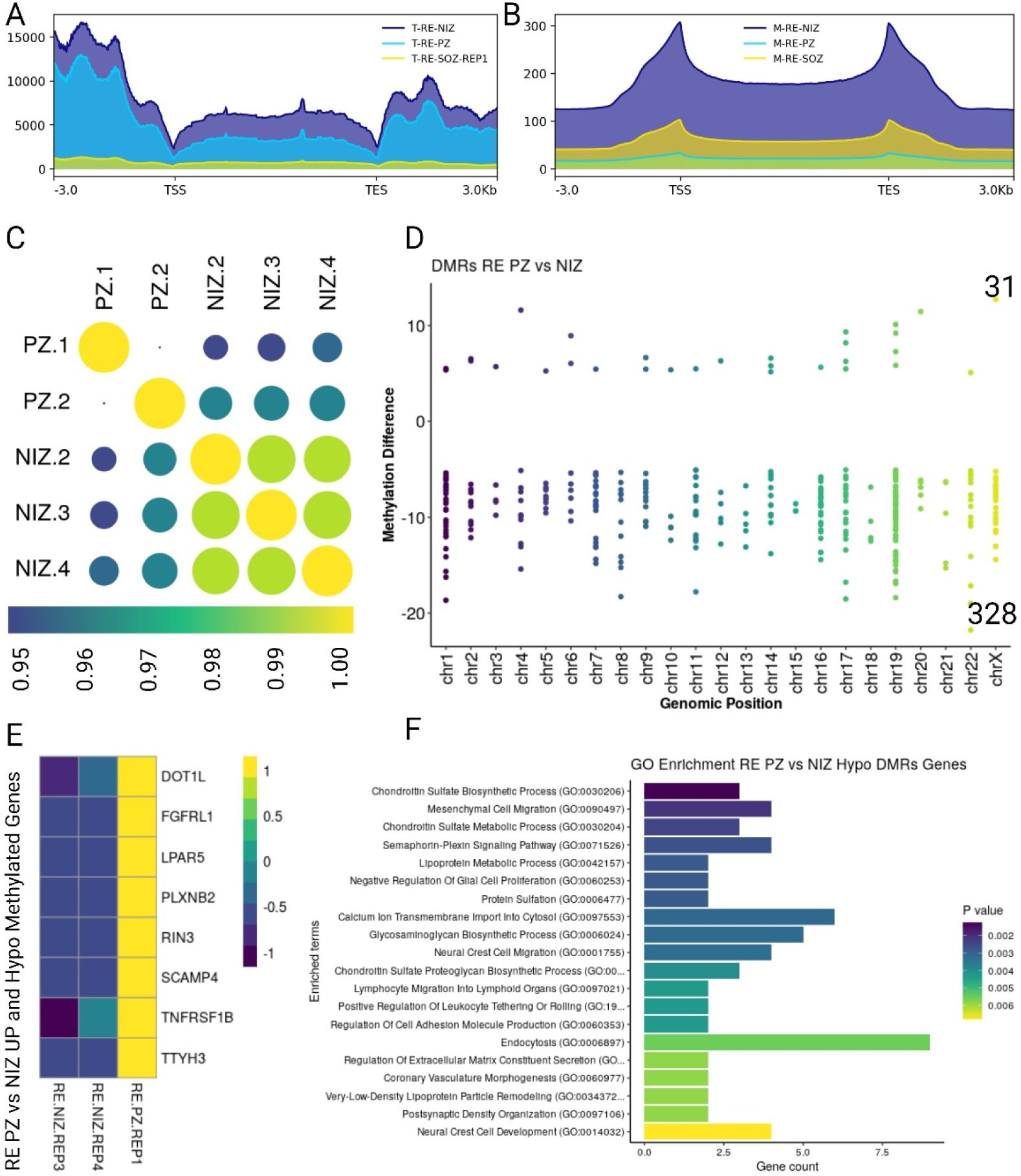
Transcriptome and methylome landscape in SOZ and NIZ regions in RE. (A-B) Transcriptome and methylome density plot in gene body and flanking regions. (C) Replicated wise correlation plot of PZ and NIZ regions methylome in RE brain. (D) A comparable number of hypermethylated (31) and hypomethylated (328) DMRs are identified when comparing the PZ to the NIZ region. (E) Up regulated and hypo methylated genes in PZ as compared to NIZ regions. (F) GO enrichment of hypomethylated and hypermethylated genes found in PZ regions.

**Figure S11.**
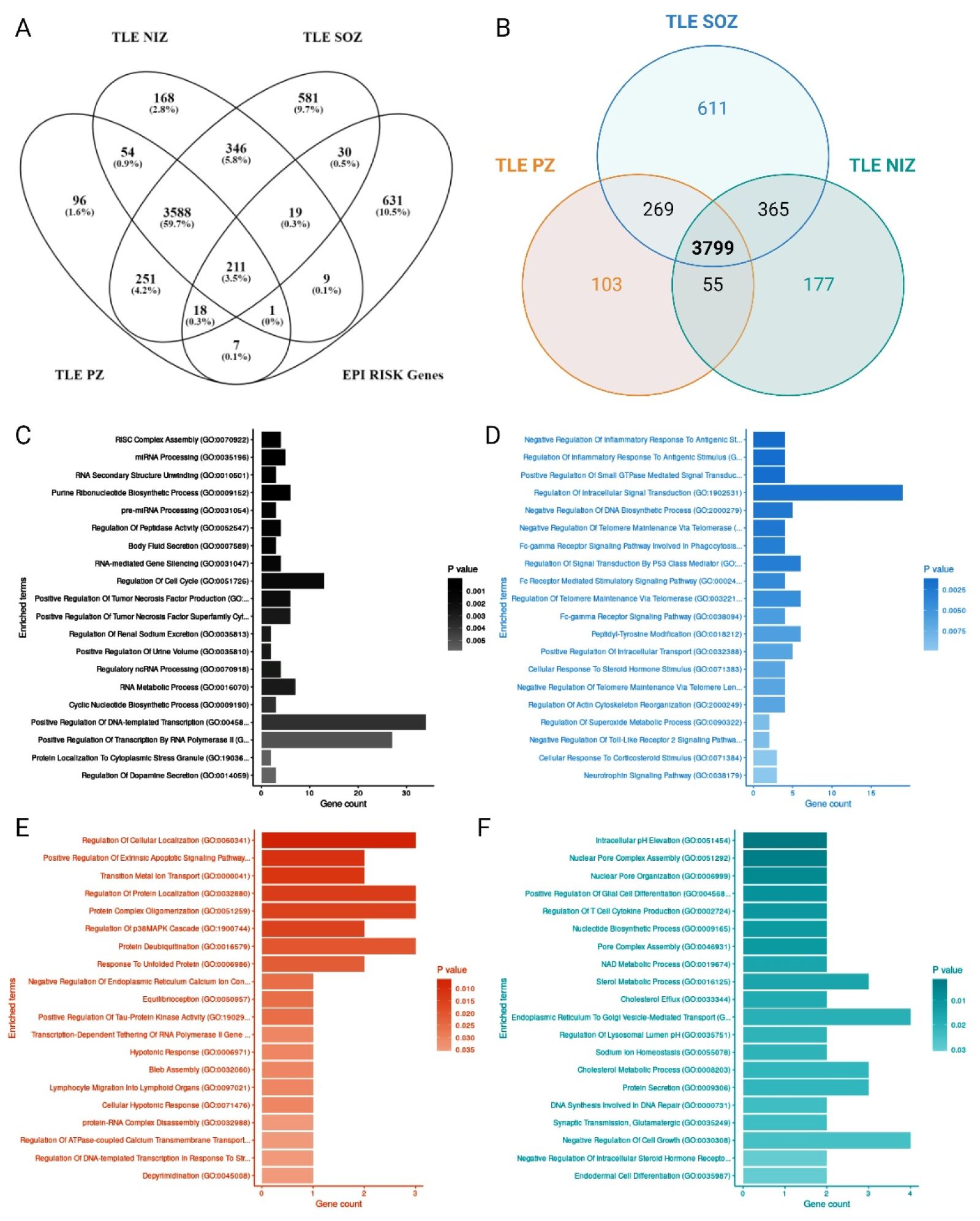
Functional enrichment of variants identified in NIZ, PZ and SOZ regions of TLE. (A-B) Venn diagram showing the overlap of genes harbouring variants in NIZ, PZ and SOZ regions of TLE brains and epilepsy risk genes, respectively. (C) GO enrichment of genes harbouring variants in all the NIZ, PZ and SOZ regions. GO enrichment of genes harbouring variants only in (D) SOZ , (E) PZ and (F) NIZ regions.

**Figure S12.**
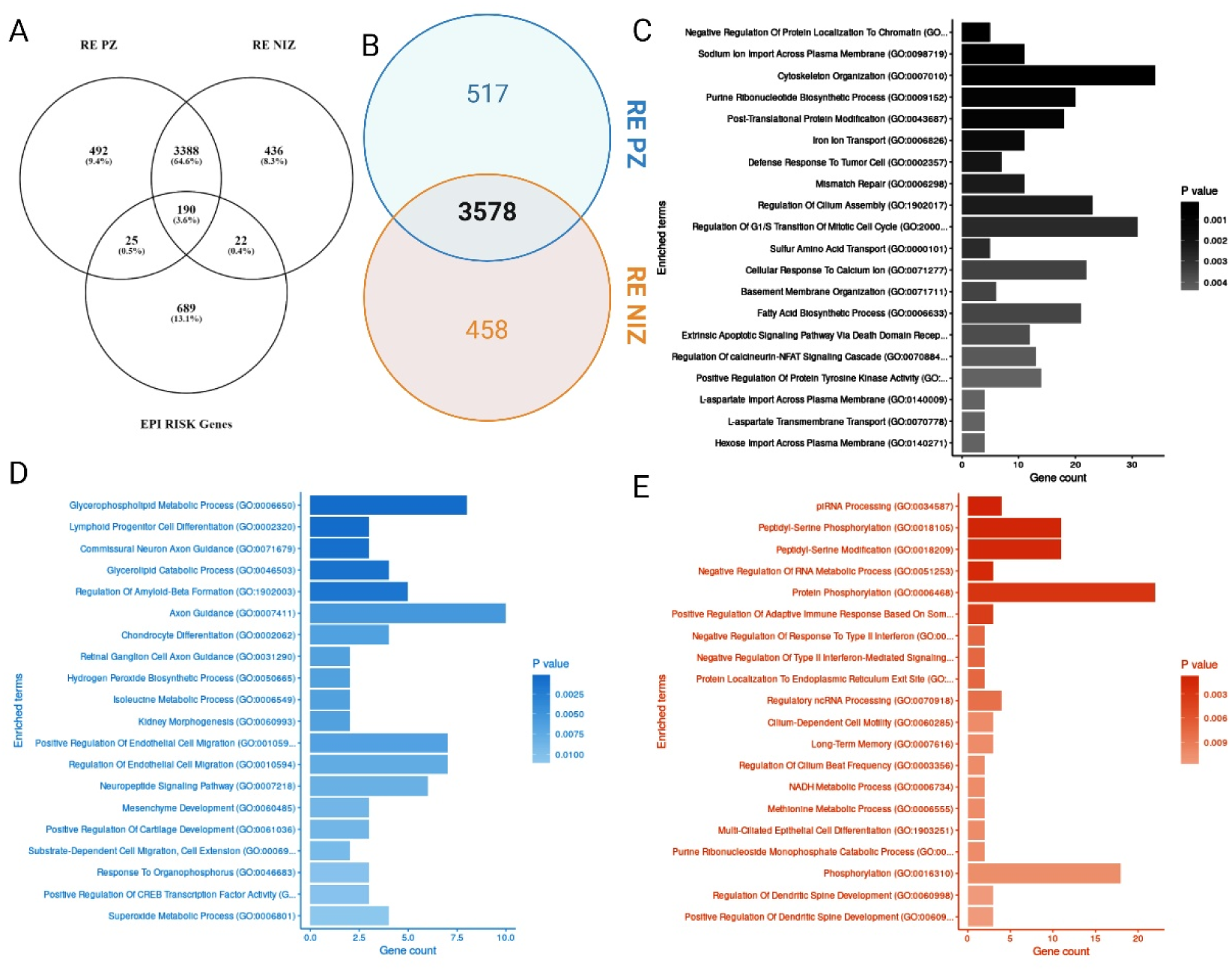
Functional enrichment of variants identified in NIZ and PZ regions of RE. (A-B) Venn diagram showing the overlap of genes harbouring variants in NIZ and PZ regions of RE brains and epilepsy risk genes, respectively. (C) GO enrichment of genes harbouring variants in all the NIZ and PZ regions. GO enrichment of genes harbouring variants only in (D) PZ and (E) NIZ regions.

## SUPPLEMENTARY TABLES

Table S1. Up regulated genes in FCD PZ vs NIZ comparison.

Table S2. Down regulated genes in FCD PZ vs NIZ comparison.

Table S3. Up regulated genes in TLE PZ vs NIZ comparison.

Table S4. Down regulated genes in TLE PZ vs NIZ comparison.

Table S5. Up regulated genes in TLE SOZ vs NIZ comparison.

Table S6. Down regulated genes in TLE SOZ vs NIZ comparison.

Table S7. Up regulated genes in TLE SOZ vs PZ comparison.

Table S8. Down regulated genes in TLE SOZ vs PZ comparison.

Table S9. Up regulated genes in RE PZ vs NIZ comparison.

Table S10. Down regulated genes in RE PZ vs NIZ comparison.

Table S11. Up regulated genes in RE SOZ vs NIZ comparison.

Table S12. Down regulated genes in RE SOZ vs NIZ comparison.

Table S13. Up regulated genes in RE SOZ vs PZ comparison.

Table S14. Down regulated genes in RE SOZ vs PZ comparison.

Table S15. Enriched GO terms in up regulated genes in FCD PZ vs NIZ comparison.

Table S16. Enriched GO terms in down regulated genes in FCD PZ vs NIZ comparison.

Table S17. Enriched GO terms in up regulated genes in TLE PZ vs NIZ comparison.

Table S18. Enriched GO terms in down regulated genes in TLE PZ vs NIZ comparison.

Table S19. Enriched GO terms in up regulated genes in TLE SOZ vs NIZ comparison.

Table S20. Enriched GO terms in down regulated genes in TLE SOZ vs NIZ comparison.

Table S21. Enriched GO terms in up regulated genes in TLE SOZ vs PZ comparison.

Table S22. Enriched GO terms in down regulated genes in TLE SOZ vs PZ comparison.

Table S23. Enriched GO terms in up regulated genes in RE PZ vs NIZ comparison.

Table S24. Enriched GO terms in down regulated genes in RE PZ vs NIZ comparison.

Table S25. Enriched GO terms in up regulated genes in RE SOZ vs NIZ comparison.

Table S26. Enriched GO terms in down regulated genes in RE SOZ vs NIZ comparison.

Table S27. Enriched GO terms in up regulated genes in RE SOZ vs PZ comparison.

Table S28. Enriched GO terms in down regulated genes in RE SOZ vs PZ comparison.

